# Glutamine Synthetase-1 induces autophagy-lysosomal degradation of huntingtin aggregates and ameliorates animal motility in a *Drosophila* model for Huntington’s disease

**DOI:** 10.1101/618629

**Authors:** Luisa Vernizzi, Chiara Paiardi, Giusimaria Licata, Teresa Vitali, Stefania Santarelli, Martino Raneli, Vera Manelli, Manuela Rizzetto, Mariarosa Gioria, Maria E. Pasini, Daniela Grifoni, Maria A. Vanoni, Cinzia Gellera, Franco Taroni, Paola Bellosta

## Abstract

Glutamine Synthetase1 (GS1) is an enzyme that catalyzes the ATP-dependent synthesis of L-glutamine from L-glutamate and ammonia as a key element of the glutamate glutamine cycle, a complex physiological process occurring between glia and neurons, necessary to control the homeostasis of glutamate. Using a *Drosophila* model for Huntington’s disease, we report that expression of GS1 in neurons ameliorates the motility defects of animals expressing the mutant *Httex1-Q93* form of the *huntingtin* gene. At the cellular level, expression of GS1 increases the basal level of autophagy and significantly reduces the size of the toxic Htt-Q93 protein aggregates. In addition, we found that expression of GS1 prevents TOR localization at the lysosomal membrane and reduction in the phosphorylation of its effector S6K. This study reveals a novel function for GS1 in neurons linking its activity to the inhibition of TOR signaling and autophagy. The identification of novel pharmacological regulators of autophagy is of particular interest considering its beneficial role in controlling neuronal health and counteracting the detrimental effects of toxic aggregates of proteinopathies including Huntington’s disease.

## Introduction

Many efforts have been focused to develop therapies to improve neuronal health, particularly those that promote autophagy, a self-cleaning process known to improve the symptoms of neurodegenerative disorders (NDDs) [1], especially for proteinopathies such as Huntington’s disease (HD) where autophagy favors the clearance of the toxic protein aggregates [2]. HD is an inherited adult-onset neurodegenerative disease characterized by the presence of an abnormal long sequence of CAG repeats in the first exon of the *huntingtin* gene (*HTT*, Chr. 4p16.3 MIM 143100), encoding a protein with a non-physiological long poly-Q trait at the N-terminus [3, 4]. HD is characterized by the progressive degeneration of neurons in the striatum and cortex. This results in motor defects, cognitive decline and death. The age of clinical onset in HD is highly variable (with a mean of ~45 years), but is strongly influenced by the length of the CAG trinucleotide expansion within the *HTT* gene. Although many cellular and molecular mechanisms are identified in HD, at present, there are no effective therapies [5, 6], and one emerging strategy to ameliorate HD is the inhibition of the TOR signaling pathway which then allows the induction of autophagy [7].

Neuronal health is also controlled by a non-autonomous cycle, called Glutamate-Glutamine Cycle (GGC) that is responsible for the exchange of glutamine and glutamate between glia and neurons thus mantaining their concentration at physiological levels [8]. Among the key enzymes controlling this cycle there are Glutamine Synthetase 1 (GS1) and Glutamate Dehydrogenase (GDH). In humans, GS1 is encoded by the *glutamate ammonia-ligase* gene (*GLUL)* that converts glutamate and ATP to glutamine, and *GLUD* that encodes GDH that catalyzes the reversible conversion of glutamate to alpha-ketoglutarate and ammonia. [9]. In humans, mutations in the *GLUL* gene are associated with a severe autosomal recessive disorder resulting in encephalopathy, organ failure and severe birth defects, probably due to the inability to detoxify ammonia [10]. *Drosophila* contains two distinct genes encoding *glutamine synthetase: GS1* and *GS2*, with *GS1* being 70.5% homologous to human *GLUL* [11] and causing embryonic lethality when mutated [12].

While the positive role of autophagy in the pathophysiology of HD is evident [2, 13], much less is known about the role of the GGC in this process. Evidences supporting the relevance of these enzymes in HD show that GS1 activity is reduced in brains from HD patients [14–17], while gain of function of GDH mutations were shown in rare polymorphisms, to favor the onset of NDDs including Parkinson and HD [18].

In order to investigate the contribute of GS1 to HD, we induced its expression using a *Drosophila* model for HD, where the expression in neurons of exon 1 of the human *HTT* gene containing 93 CAG repeats (hereon referred to as *Httex1-Q93)* recapitulates some of the cellular and molecular events described in HD patients, including loss of neurons and animal motility [19]. Our experiments show that the ectopic expression in neurons of a functional *GS1* enzyme, along with *Httex1-Q93*, significantly improves animal motility and rescues neuronal loss caused by *Httex1-Q93* expression. At the cellular level, we found that expression of GS1 considerably decreases the size of Htt-Q93 protein aggregates in neurons, a process that is accompanied by an increase in basal level of autophagy. Autophagy is a self-eating cellular mechanism normally induced when nutrients are reduced, which is counteracted by TOR signaling. Activation of TOR starts with the formation of the RagA/B- RagC/D GTPases heterodimer complex, stimulated by amino acids [20] that together with the GTPase Rheb [21–23] recruits TOR at the lysosomal membrane to form the TORC1 complex. Activation of TOR culminates with the phosphorylation of its targets proteins S6K and 4EBP [24, 25]. Interestingly, we found that co-expression of GS1 together with *Httex1-Q93* partially reduced S6K phosphorylation, suggesting that expression of GS1 reduces TORC1 activity. Amino acid analysis from the heads of animals expressing GS1 in neurons reveals an increase in glutamine levels but also a significant decrease of few essential amino acids, such as histidine, proline, isoleucine, and of arginine. Our data suggest that expression of GS1 induces a condition that simulate “amino acid starvation” in neurons, which is necessary to induce autophagy to replenish the pool of amino acids that are reduced. In addition, we found an opposite regulation of aspartate with respect to asparagine, which increases in the presence of GS1 suggesting an up-regulation of the activity of the enzyme asparagine synthetase (ASNS) that converts aspartate into asparagine from glutamate with the use of glutamine. Interestingly, ASNS was shown to be necessary for the regulation of the uptake of amino acids, especially serine, arginine and histidine, essential for the survival of tumor cells [26, 27]. Overall, our data suggest that the expression of GS1 in neurons results in a reduction of essential amino acids that in turn results in autophagy. Glutamine levels are not reduced but perhaps used, together with aspartic acid, in asparagine catabolism, to constitute a feedback loop that maintain autophagy active and favors cellular survival. In conclusion, we show a novel role for GS1 that induces autophagy and reduces the formation of Htt-Q93 toxic aggregates to ameliorate the motility defects in a *Drosophila* model for HD. Understanding the mechanisms that link the metabolic pathways activated by GS1 to amino acids signaling in neurons, would be an initial step to delineate a novel function for this enzyme, part of the GGC, in the brain necessary to control autophagy and to induce neuronal survival not only in pathological situations such as HD but also in physiological conditions during neuronal homeostasis and in ageing.

## Results

### *GS1* rescues neuronal death induced by expression of *Httex1-Q93* in the retina

GS1 expression was found reduced in post-mortem brains from HD patients [14–17]. In addition, human fibroblasts from HD patients have increased levels of glutamate that correlate with (CAG)_n_ length in the *HTT* gene (Fig. 1A). Since GS1 converts glutamate into glutamine, we investigated whether overexpression of *Drosophila* GS1 in neurons could reduce glutamate concentration and ameliorate the pathological effect of mutant *Htt* (*mHtt*) in a *Drosophila* model for HD generated by expressing *Httex1-Q93*. This model was shown to recapitulate many of the cellular and molecular events described in neurons of human HD patients including motility defects [28]. In *Drosophila,* two distinct genes (*GS1* and *GS2*) encode glutamine synthetase [12]. *GS1* (Fig. 1B) is 70.5% homologous to human *GLUL*/*GS1* [11] (Supplementary Fig. 1A) and is the prevalent form expressed in neurons [12]. Expression of *GS1 in vivo*, using the ubiquitous *actin-Gal4* promoter, results in the production of a *GS1* mRNA (Supplementary Fig. 1B) encoding a 50 kDa protein (Fig. 1C) which correlates with a specific increase of GS1 enzyme activity (Fig. 1D). We first analyzed in the retina, using the tissue-specific *GMR-Gal4* promoter, whether co-expression of *GS1* and *Httex1-Q93* was able to reduce neuronal degeneration of eye pigment cells, as previously shown in the *Httex1-Q93* HD model [28, 29]. In our experiments, co-expression of *GS1* and *Httex1-Q93* completely rescued the lack of pigmentation in the ommatidia induced in neuronal cells by *Httex1-Q93* expression (compare Fig. 1F with 1G). Expression of GS1 alone did not alter eye morphology and pigmentation was similar to that of control *Httex1-Q16* animals (Figs. 1H and 1E). On the contrary, reduction of GS1 using RNA interference (RNAi) exacerbated the defects in the ommatidia of animals expressing *Httex1-Q93* and these animals showed premature loss of pigment cells that at 8 days after eclosion (DAE) was similar to that of animals expressing *Httex1-Q93* at 20 DAE (Supplementary Fig. 2A). A detailed ultramicroscopic analysis of the tissues underneath the ommatidia from flies expressing *Httex1-Q93* showed a remarkable alteration of cornea morphology with the presence of large vacuoles indicating loss of the cells underneath the retina (Fig. 1J). Expression of *GS1* ameliorates this phenotype and also restricted cell death to a small area beneath the cornea (Fig. 1K). Morphological analysis of the retina from *GS1* and control *Httex1-Q16* animals revealed no defects (Fig. 1L and I, respectively).

**Fig 1:**
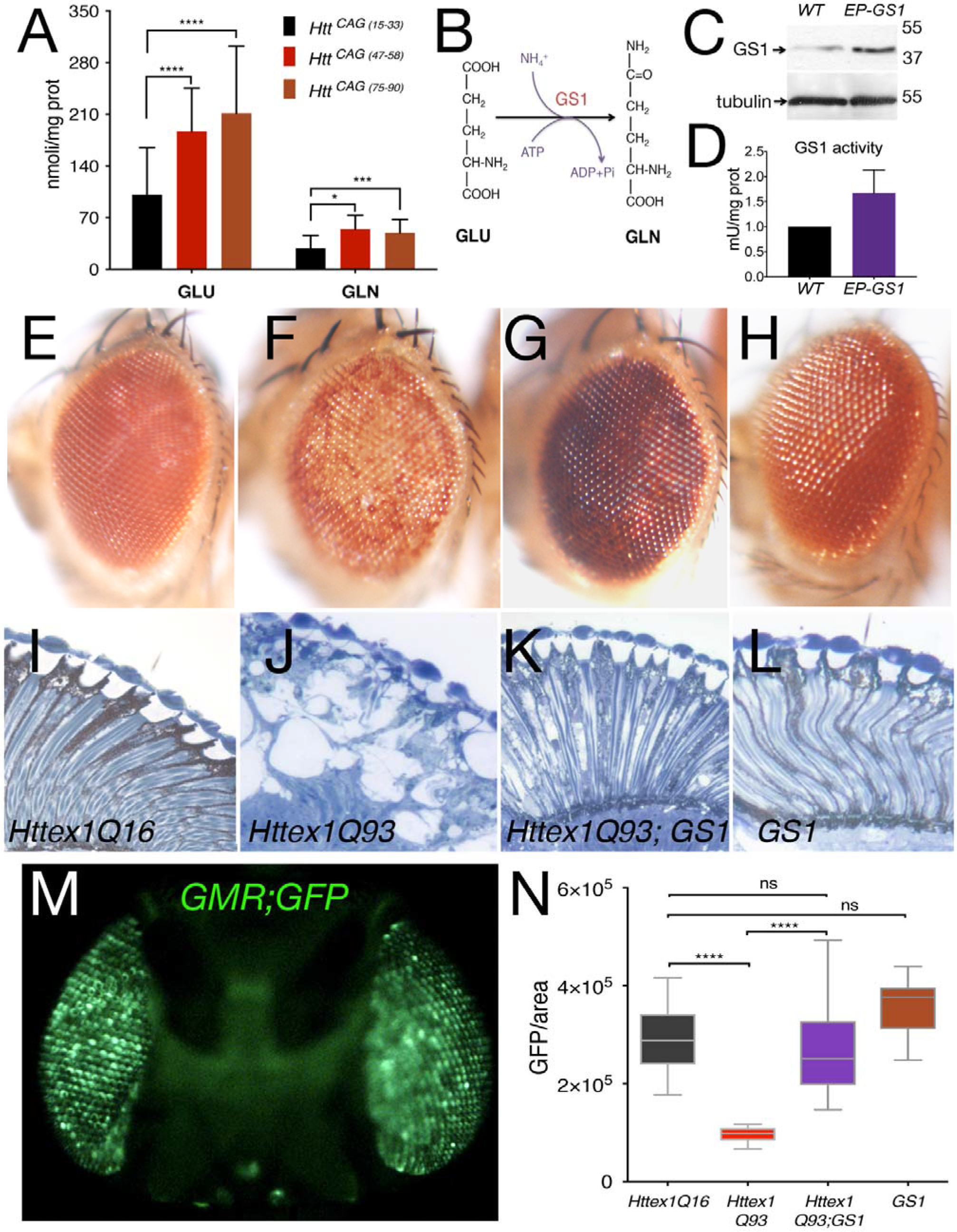
Expression of GS1 in the retina rescues neuronal death induced by Htt-Q93. A) Quantification of glutamate and glutamine (MS/HPLC) in cultures of human fibroblasts from control donors or patients carrying different length of the CAG trait in the *HTT* gene. (B) Schematic chemical reactions controlled by the Glutamine Synthetase 1 enzyme. (C) Western blot from larvae showing the relative amount of GS1 protein from *wt* animals or expressing the EP {GS1} transgene using the *actin-Gal4* promoter. Tubulin was used as loading control. (D) GS1 enzymatic activity in extracts from C. (E-H) Photographs of *Drosophila*’s compound eyes (lateral view) from females at 10 days after eclosion (DAE) expressing the indicated *UAS*-transgenes using the *GMR-Gal4* driver. (I-L) Representative photographs of retinal sections from animals as in E-H. (M) Fluorescent image of the head of *GMR-Gal4; UAS-GFP* animals (sagittal view). (N) Quantification of GFP from photographs of adult eyes (see methods). The * = *p* < 0.05, *** = *p* < 0.001, **** = *p* < 0.0001 values in panels A and N, were calculated from Student’s *t*-test from at least three independent experiments, error bars indicate the standard deviations.

Neuronal degeneration can be analyzed by measuring the decay over time of the intensity of GFP expressed in cells of the retina [29]. *GFP* was expressed in the retina together with our transgenes and GFP intensity analyzed (Fig. 1M). These experiments showed that expression of *Httex1-Q93* significantly decreases GFP fluorescence in the ommatidia quantified at 20 DAE (ANOVA *P*<0.0001) (Fig. 1N). Moreover, co-expression of *GS1* completely rescued this defect supporting the hypothesis that GS1 activity counteracts neuronal death induced by *Httex1-Q93* expression in the retina cells. GFP intensity in control *GMR-Q16* and *GS1* animals (Fig. 1N) was not significantly different (ANOVA *P*=*ns*). (ANOVA data-analysis is in the supplementary Excel file).

### *GS1* expression in neurons improves locomotion defects of *Httex1-Q93* animals

To analyze if the ability of GS1 to rescue neuronal degeneration in the retina was extended to neurons controlling animal motility, we co-expressed *GS1* and *Httex1-Q93* using the *elav*^*c155*^-*Gal4* pan-neuronal promoter, hereon referred to as *elav-Gal4* [28]. Analysis in larval brains for proper expression of the transgenes showed that *Httex1-Q16* and *Httex1-Q93* encode a 35 kDa and a 55 kDa protein, respectively. Expression of *Httex1-Q93* also results in the formation of high-molecular-weight bands (HMW-mHtt) entrapped in the stacking gel and visible also in brains co-expressing *Httex1-Q93* and *GS1* (Fig. 2A). The presence of these high molecular weight forms reacting with the anti-Htt antibody is a characteristic pattern of expression for the mutant Htt, as previously showed in *Drosophila* heads expressing the same *Httex1-Q93* transgene [29, 30].

**Fig 2:**
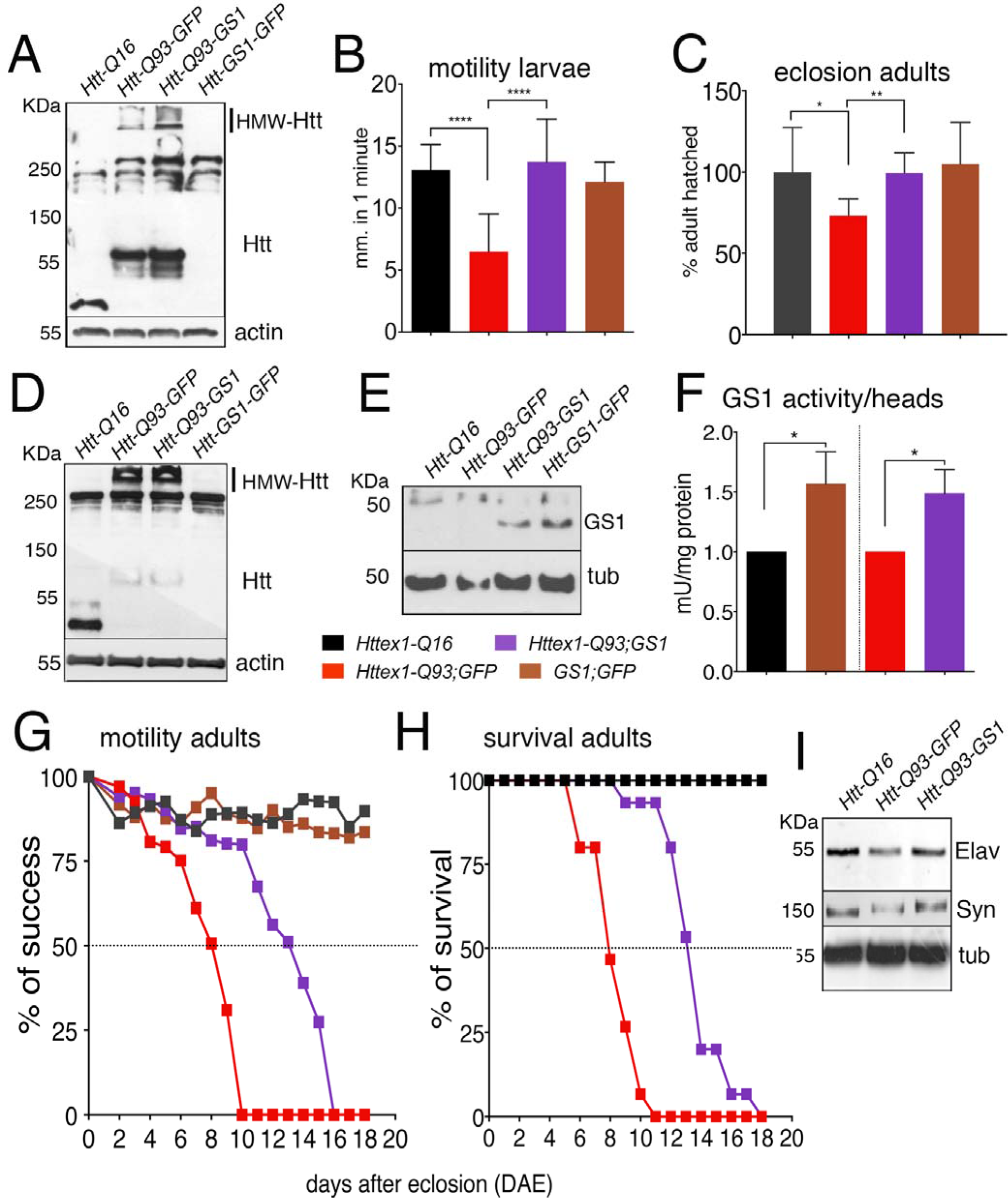
Expression of GS1 in neurons rescues motility defects and neuronal death induced by Htt-Q93. (A) Western blot showing the relative amounts of Htt protein in lysates from 3rd instar larval brains expressing the indicated transgenes using the pan-neuronal driver *elav-Gal4*, Actin was used as loading control. The dark line on the right indicates the stacking gel. (B) Quantification of larval motility of 3rd instar larvae of the indicated genotype. (C) Adult eclosion-timing of animals. (D-E) Western blots showing the relative amounts of Htt and GS1 proteins in lysates from heads of females at 8 days after eclosion (DAE) expressing the indicated transgenes using the *elav-Gal4* driver, Actin was used as control. (F) GS1 enzymatic activity in lysates from heads of females at 8 DAE. (G-H) Quantification of motility in females (G) and relative % of survival (H) of adult animals expressing the relative transgenes in neurons. (I) Western blot showing the amounts of endogenous neuronal specific proteins elav and synapsin in lysates from the heads of animals of the indicated genotype, Tubulin was used as control. The * = *p* < 0.05, *** = *p* < 0.001, **** = *p* < 0.0001 values in panels B, C, F, G were calculated from Student’s *t*-test from at least three independent experiments, error bars indicate the standard deviations. One-way ANOVA analysis of graph G and H are in the supplementary Excel file.

Since one hallmark of HD is a progressive impairment in the movement [28], we investigated whether expression of *GS1* could rescue animal motility defects induced by *Httex1*-Q93 expression in neurons. We first analyzed differences in larval motility by comparing the ability of third instar animals to crawl in an arena (Supplementary Fig. 3C-E, movies). These experiments showed that the motility defects induced by *Httex1-Q93*, visible already at 48-72 hours after egg laying (AEL) (Fig. 2B, ANOVA *P*<0.0001), are completely rescued by co-expression of *GS1* (ANOVA *P*=ns). Adults *Httex1-Q93* hatched at the expected developmental time without any visible morphological defects but with a 30% of lethality at the stage of pharates (Fig. 2C). Co-expression of *GS1* completely rescued also the hatching defects, while animals expressing *GS1* alone did not show any visible defect in motility (Fig 2B, ANOVA ns), and have a survival rate similar to control *Httex1-Q16* (Fig 2C). In adult animals, western blot analysis using lysates from the heads of females at 8 DAE show a pattern of expression for Htt-Q16 and Htt-Q93 proteins similar to that of the larval brains (Fig. 2D). GS1 protein levels and enzyme activity were similar in the heads of flies expressing *GS1* alone or co-expressing *Httex1-Q93* (Fig. 2E). Moreover, GS1 enzyme activity was inhibited with the same efficiency in both genotypes by the specific GS1 inhibitor methionine sulfoximine (MSO) (Supplementary Fig. 4). To analyze if co-expression of GS1 ameliorates the motility defects of adult flies, we performed climbing assays over a period of time (Supplementary Fig. 3F, movie). These data show that *elav*/+*; Httex1-Q93*/+ females have a significantly reduced climbing activity (50% reduction at 8 DAE) as compared to that of *Httex1-Q16* control flies (ANOVA *P*<0.0001). Co-expression of *GS1* ameliorates these defects and *elav*/+*; Httex1-Q93*, *GS1*/+ females showed a delayed reduction in their motility but (50% at 12 DAE) (Fig. 2 G, ANOVA *P*<0.05). These results were confirmed using three independent *Httex1-Q93*, *GS1* recombinant lines (Supplementary Fig. 5). Motility reduction is often accompanied by a decrease in the survival rate of the animals (Fig. 2H). Expression of *GS1* was also able to reduce lethality of *elav*/+*; Httex1-Q93*/+ flies (Fig. 2H, ANOVA *P*<0.05), while flies expressing *GS1* alone and control *Httex1-Q16* did not show any visible alterations in their motility or viability.

In males, expression of *Httex1-Q93* revealed a very low survival rate and *elav*/*Y; Httex1-Q93*/+ animals have defects in climbing activity that was 50% reduced at about 2-3 DAE (Supplementary Fig. 6A-B). The stronger effect of *Httex1-Q93* in males is probably due to a dosage compensation effect since the *elav-Gal4* transgene is inserted in the *X* chromosome [31].

To estimate the rescue of neuronal death by GS1, we analyzed the expression of neural markers in lysates from the heads of females at 8 DAE. These experiments showed that co-expression of *GS1* rescues the reduction of elav and synapsin protein levels, detected in cell extracts from animals expressing *Httex1-Q93* alone (Fig. 2I), suggesting that the activity of GS1 rescues neuronal death in the adult heads

### Expression of *GS1* reduces Htt-Q93 protein aggregates in neurons

Mutant Htt forms large toxic intracellular aggregates. Expression of *Httex1-Q93* using the *elav* promoter results in large protein aggregates clearly visible by immunofluorescence in the gigantic cells of *Drosophila* larval salivary glands (Fig. 3A) where the *elav-Gal4* promoter is also transcriptionally active. Since co-expression of *GS1* restored the motility defects of *Httex1-Q93* animals, we analyzed if its activity could result in the reduction of the size of Htt-Q93 aggregates. Aggregates in neurons were visualized by immunofluorescence, using anti-human Htt antibodies that clearly detected the presence of Htt-Q93 aggregates in cells from the calyx already at 72-90 hours of development, but not in neurons of control *Httex1-Q16* animals (Fig. 3B-C). We then focused our analysis on the Intra-Lobe (IL), and the Ventral Nerve Cord (VNC) regions of the larval brain. This analysis show that co-expression of *GS1* significantly reduced the size of Htt-Q93 aggregates in both IL (Fig. 3E-H, *t*-student, *p*<0.01), and VNC (Fig. 3F-I, *t*-student, *p*<0.01). These data were confirmed by filter assays that trap the larger aggregates in the lysates from larval brains, into a membrane of nitrocellulose (Fig 3G). No aggregates were detected in brains from animals expressing *Httex1-Q16* or *GS1* alone. Similar results were obtained in adults, where the expression of *GS1* significantly reduced the size of Htt-Q93 aggregates, measured by immunofluorescence analysis in the heads of females at 8 DAE (Fig. 3K-L, *t*-student, *p*<0.001), and also visualized by filter trap assays (Fig. 3M). Altogether, our data suggest that GS1 induces a mechanism that helps or favors the degradation of the toxic mHtt-Q93.

**Fig 3:**
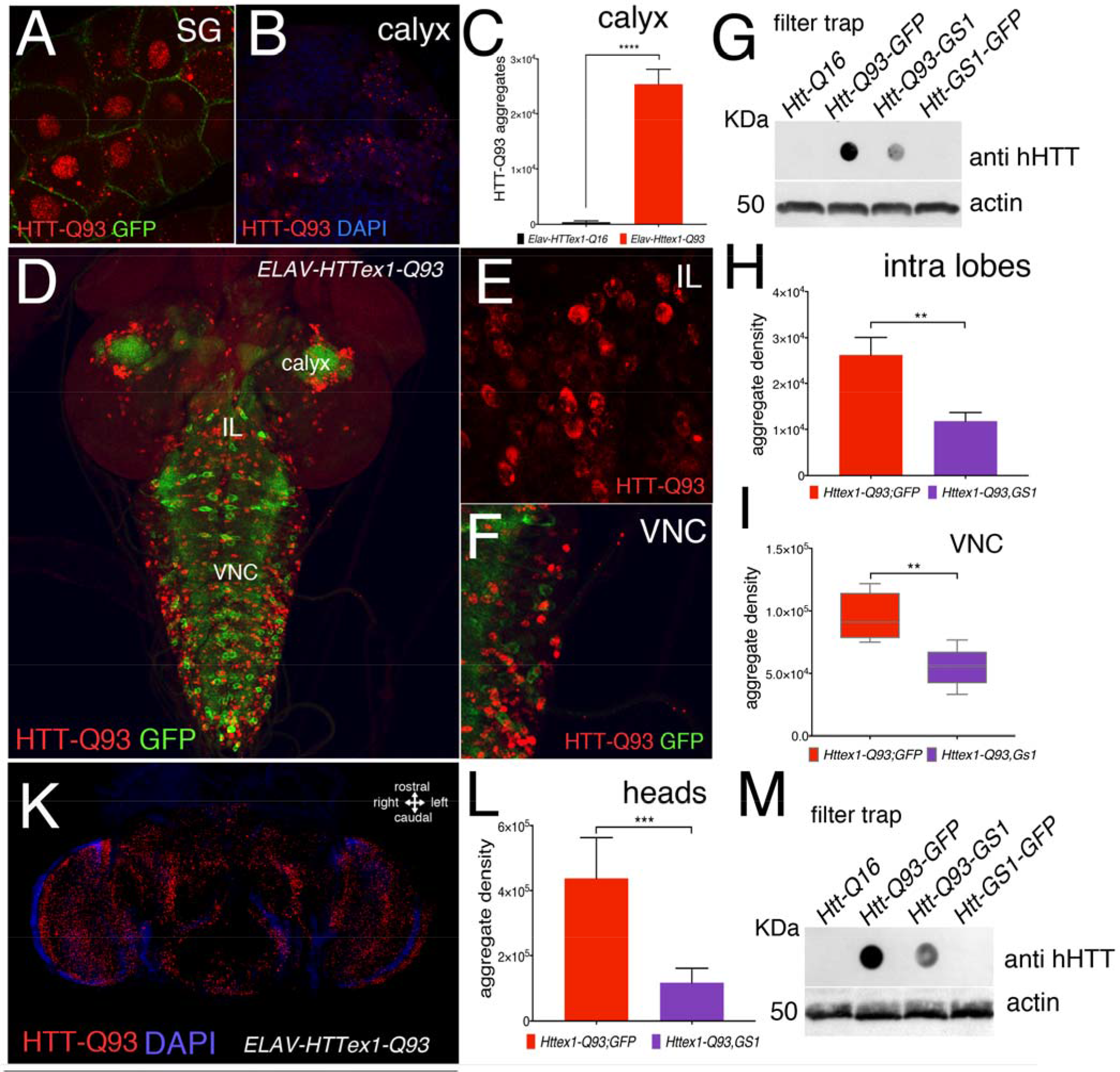
Expression of GS1 in neurons reduces the size of Htt-Q93 aggregates. Confocal photographs showing Htt-Q93 protein aggregates in the cytoplasm and surrounding the nuclei of cells in the salivary glands (SA) where the *elav-Gal4* insertion is also transcriptionally active. Aggregates were visualized by immunofluorescence using anti human Htt (hHtt) antiserum. Htt-Q93 aggregates in neurons from the region of the brain called calyx. (C) Quantification of the Htt-Q93 aggregates in the region of the calyx of *Httex1-Q93* or *Httex1-Q16* control animals; aggregate density was calculated from the integrated density of the immunofluorescence signal using anti hHtt antibody from 10 confocal *z*-stacks each animal and elaborated with *imageJ*, as explained in Materials and Methods. (D) Confocal image of the whole larval brain of the genotype *elav-Gal4; Httex1-Q93; mGFP* that marks neurons with mGFP, while the Htt-Q93 aggregates are visualized by immunofluorescence using anti hHtt antiserum. (E-F) Htt-Q93 aggregates visualized in the Intra-Lobes (IL) and in the Ventral Nerve Cord (VNC) region respectively. (G) Filter trap analysis using lysates from the brains of the indicated transgenes (upper panel); the expression of Actin was analyzed by western blot in the same samples, as internal control for loading (lower panel). (H-I) Relative quantification of Htt-Q93 protein aggregates in the IL and VNC regions. (K) Confocal image of *Drosophila elav-Gal4; Httex1-Q93* adult female’s head at 10 DAE, immune-stained for the presence of Htt-Q93 aggregates (RED, nuclei DAPI). (L) Quantification of the Htt-Q93 aggregates in the heads of the animals of the indicated genotype. (M) Filter trap analysis of Htt-Q93 protein aggregates using lysates from the heads (upper panel). Actin was used as the internal control (lower panel). The ** = *p* < 0.01, *** = *p* < 0.001 values in panels C, H, I, L, were calculated from Student’s *t*-test from at least three independent experiments, error bars indicate the standard deviations.

### GS1 induces autophagy in neurons expressing Httex1-Q93

In neurons, one of the major pathway for the clearance of toxic aggregates is the autophagy-lysosomal system [32]. To investigate whether GS1 may activate an autophagy-lysosomal pathway in neurons, which may explain the reduction of Htt-Q93 aggregates, we analyzed the formation of the characteristic autophagosome vesicles *in vivo* using the reporter *mCherry-Atg8a*. This construct expresses the autophagosome-associated ubiquitin-like protein Atg8a fused with the fluorescent mCherry protein allowing the visualization of autophagosomes as fluorescent punctae/vesicles visible under fluorescence microscope [33]. To identify neurons, we co-expressed *mGFP* under the control of the *elav-Gal4* promoter (Fig. 4C). Quantification of the intensity of fluorescence of *mCherry-Atg8a* in the autophagosome, was analyzed in the region of the calyx (Fig. 4A, B, and D) and in the Ventral Neuronal Cord (VNC) (Fig. 4C-E) from 3rd instar larval brains. This analysis showed that co-expression of *Httex1-Q93* and *GS1* significantly increased the formation of autophagosomes in the calyx neurons (Fig. 4D) (Student *t*-test, *p*<0.0001) and in the VNC (Fig. 4E) (Student *t-*test, *p*<0.05). Expression of GS1 alone exhibited a substantial increase in the number of autophagosomes in both regions of the larval brain (Fig. 4 D-E), while expression of *Httex1-Q93* resulted in the reduction of the number of autophagosomes, particularly visible in neurons from the calyx (Fig. 4D) (Student *t*-test, *p*<0.01). A similar analysis was performed in the heads of adult females, where the intensity of the autophagosome was quantified in five different regions of the head of animals at 8 DAE (Fig. 4F-G). These experiments confirmed that co-expression of *GS1* resulted in a significant increase in the number of the autophagosomes in *Httex1-Q93* animals (Student *t*-test, *p*<*0.01*). Moreover, also in adult did the expression of *GS1* increase the number of autophagosomes compared to control *Httex1-Q16* animals (Student *t*-test, *p*<*0.05*).

We then analyzed if the ability of GS1 to ameliorate motility in *Httex1-Q93* animals was dependent on a functional autophagic pathway by genetically reducing the expression of *Atg1*/*ULk1* or *Atg5*, key components of macroautophagy, using RNAi lines [34]. These experiments showed that reducing either *Atg1* or *Atg5* completely abolished the ability of *GS1* to ameliorate the climbing defects in *Httex1-Q93; GS1* animals (Fig. 4H) (One-way-ANOVA data-analysis are in supplementary Excel file), confirming our hypothesis that the ability of GS1 to ameliorate the motility defects in HD animals depends on an active autophagic pathway. In autophagy, the size of the autophagosome is regulated by the amount of Atg8a protein recruited to the vesicles during maturation of the phagosome, and by the presence of the p62/SQSTM1 (sequestosome 1), which recruits ubiquitinated substrates to the autophagosomes, features conserved also in *Drosophila* [35]. In the autophagosome, the Atg8a protein (15 kDa) is lipidated and cleaved to yield a fast migrating band visible in western blot as an approximately 7-kDa protein. Direct interaction of p62 with Atg8a, by virtue of its ubiquitin-associated domain, results in p62 degradation. Thus, we analyzed the expression level of Atg8a and p62 proteins as markers for a functional autophagy flux. Our analysis of the adult heads, showed that expression of *GS1* increases the amount of the fast-migrating cleaved Atg8a-II form (2.7-fold) and a similar effect was observed when *GS1* was co-expressed with *Httex1-Q93* (2.1-fold) (Fig. 4I). Moreover, analysis of p62 expression revealed a reduction in the heads of *Httex1-Q93* animals co-expressing *GS1* (0.25-fold) compared to control *Httex1-Q16*, and a similar decrease was observed in animals expressing *GS1* alone (0.35-fold) (Fig. 4I). Altogether, these data suggest that GS1 activity induces an active autophagic flux also in neurons from animals carrying mHtt.

**Fig 4:**
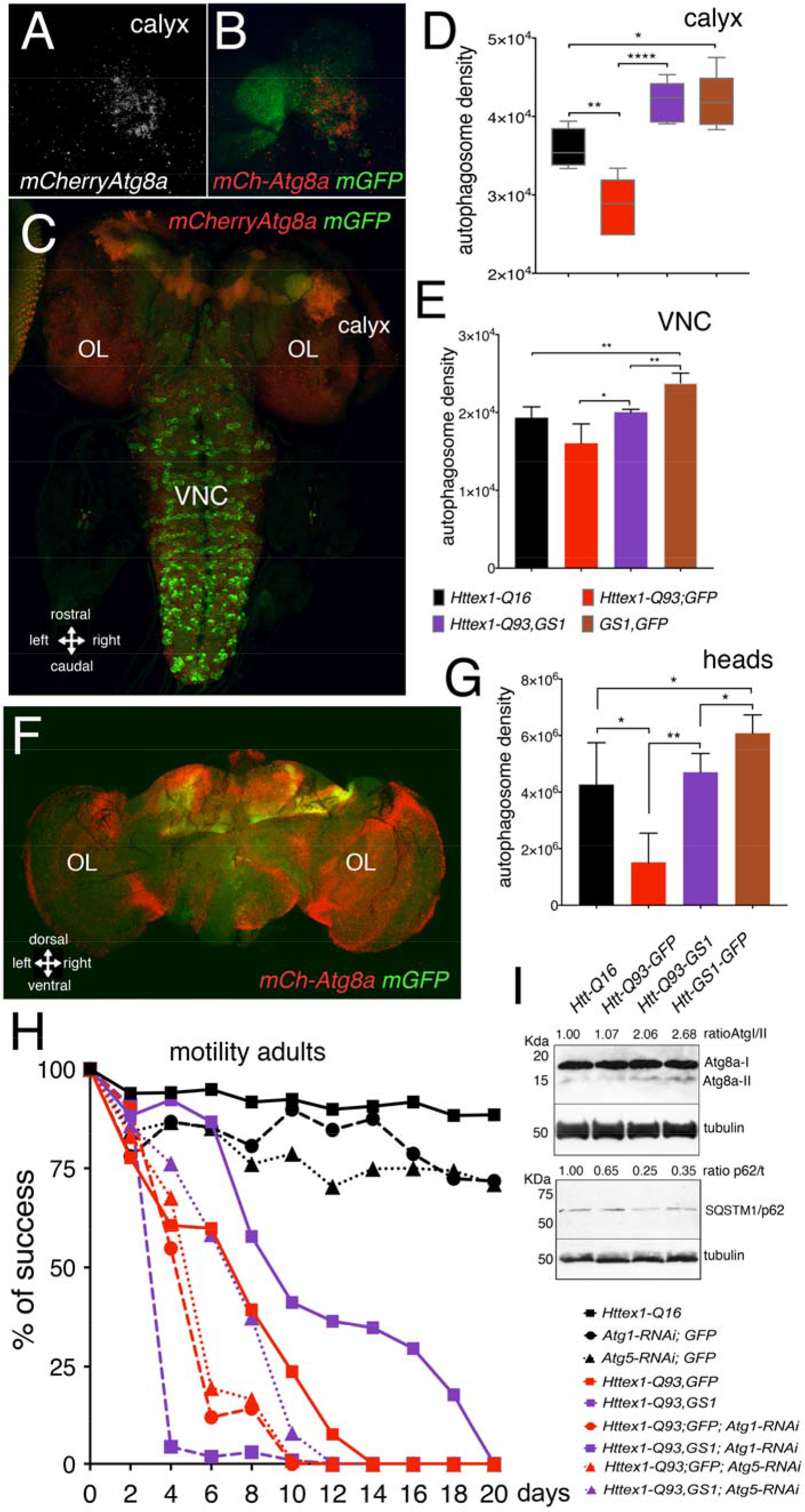
Expression of GS1 in neurons requires a functional autophagy to rescue the motility defects induced by Httex1-Q93. (A-B) Photographs of the calyx region from the brain of 3^rd^ instar larvae expressing under the *elav-Gal4* promoter the *UAS-mCherry-Atg8a* as marker of autophagy and *UAS-mGFP* to marks neuronal’s membrane. (C) Photograph of a whole brain showing the pattern of expression of *elav-mCherry-Atg8a; mGFP* in 3rd instar brain. (D-E) Quantification of autophagy, measured as the integrated density and then converted in autophagosome density (see methods) of *mCherry-Atg8a* in the autophagosome from neurons of the calyx (D) and from the Ventral Nerve Cord (VNC) (E) in animals of the indicated transgenes. (F) Photograph of an adult head of the genotype *ELAV-mCherry-Atg8a; mGFP*. (G) Quantification of autophagosome density in the adult heads or the relative transgenes. (H) Motility assay in adult females showing that reduction of *Atg1* and *Atg5* expression significantly impairs the ability of GS1 to rescue the motility defects induced by *Httex1-Q93*. The efficiency of *Atg1* and *Atg5 RNAi* lines was tested in western blot experiments and in the ability to reduce autophagy, see supplementary Fig. 7. (I) Western blot using lysates from the heads of females at 10 DAE of the indicated genotype, showing the pattern of expression of Atg8a and of SQSTM1/p62 proteins as a read-out for autophagy, tubulin was used as control. The * = *p* < 0.05, ** = *p* < 0.01, *** = *p* < 0.001, **** = *p* < 0.0001 values in panels D, E and G were calculated from Student’s *t*-test from at least five animals, error bars indicate the standard deviations. One-way ANOVA analysis for panel H is in the supplementary Excel file.

### Expression of GS1 reduces S6K phosphorylation and TOR localization at the lysosomal membrane

Autophagy is inhibited by the presence of essential amino acids that stimulate TOR kinase to translocate into the lysosomal membrane and to be assembled into the TORC1 complex, resulting in its activation and phosphorylation of downstream signaling targets, including S6K and 4EBP [36]. To explore the ability of GS1 to affect TOR signaling, we analyzed the level of phosphorylation of S6K on Thr-398 in lysates from the heads of adult animals. This analysis revealed that expression of GS1 in neurons reduces the levels of S6K phosphorylation (ratio 0.42/1.00, *p*<0.05) (Fig. 5A), a condition that is conserved also when *GS1* is co-expressed with *Httex1-Q93* (ratio 0.76/1.00, *p*<0.05). To analyze whether the reduction of S6K phosphorylation was due to an impairment of TOR translocation to the lysosome, we analyzed its co-localization with the lysosome membrane marker LAMP1-GFP. TOR kinase was visualized by immunofluorescence using an antibody against human TOR, which also recognizes the *Drosophila* orthologue (Fig. 5B, and Supplementary Fig. 8). This analysis showed that TOR localization was significantly reduced in lysosomes from neurons expressing *GS1* compared to that in neurons from control *Httex1-Q16* animals, where is normalized to 1 (Fig. 5C, compare brown to black lanes, Student *t-*test, *p*<0.05). This reduction was maintained also in animals co-expressing *GS1* and *Httex1-Q93* (Fig. 5C, compare purple to black lanes, *p*<0.01). Moreover, a significant reduction in TOR localization was detected in the lysosome from neurons of animals co-expressing *GS1* with *Httex1-Q93* (Fig. 5C, compare purple to red lanes, Student *t-*test, *p*<0.05) indicating that GS1 activity is able to induce a mechanism that inhibits the displacement of TOR from the lysosome membrane also in HD animals.

**Fig 5:**
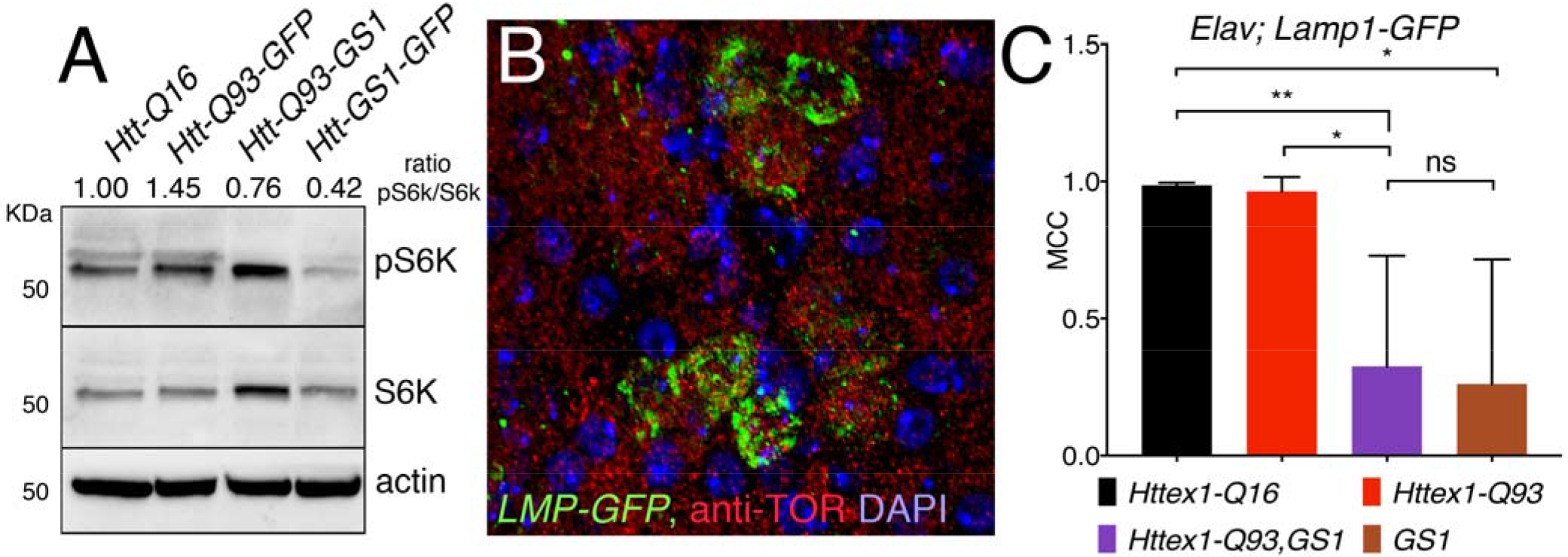
Expression of GS1 in neurons reduces TOR signaling and its localization at the lysosome membrane. (A) Western blot showing the level of phosphorylation of S6K on Thr 398, and S6K total levels in lysates from the heads of animals expressing the indicated transgenes in neurons using the *elav*- *Gal4* driver. Actin expression shows loading control. The ratio between pS6K/S6K was calculated from three independent experiments (*p*< 0.0001). (B) Photograph representing the region in the intra lobe (IL) of 3^rd^ instar larval brain that was used to analyze the co-localization between LAMP1-GFP as a marker of the lysosome membrane and TOR protein localized by immunostaining using an anti TOR antibody, (Red). (C) Quantification of the co-localization between LAMP1-GFP and TOR (Red) in the neurons from the IL region of larval brains was quantified using by *ImageJ* using the JACoP plugin and Pearson’s algorithm with Manders Coefficients and Costes automatic threshold (MCC). The * = *p* < 0.05, ** = *p* < 0.01, values in panels C were calculated from Student’s *t*-test from at least four independent animals.

### Expression of GS1 in neurons increases glutamine levels

Similar to our data in fibroblasts from HD patients (Fig. 1A), we found that glutamate concentration is increased in the heads of animals expressing *Httex1*-Q93 (Fig. 6A, solid red vs. black columns, *t-* test *p*<0.0001). Expression of GS1, which converts glutamate into glutamine, partially lowers the high concentration of glutamate while increasing the level of glutamine (Fig. 6A, purple columns vs. red columns). No significant change in glutamate level was noted between *GS1* animals and control *Httex1-Q16* (Fig. 6A solid brown vs. black columns), while glutamine levels are strongly increased (empty black vs. brown columns, *t*-test *p*<0.0001), an effect that was also detected in animals co-expressing *GS1* and *Httex1-Q93* (empty black vs. purple columns, *t-*test *p*<0.0001). In addition, we also found a significant increase in glutamine levels in the heads of *Httex1-Q93*; GS1 animals compared to *Httex1-Q93* alone (empty purple and red lanes, *t*-test *p*<0.05), suggesting that GS1 *in vivo* favors the conversion of glutamate into glutamine also in the presence of *Httex1-Q93*.

**Fig 6:**
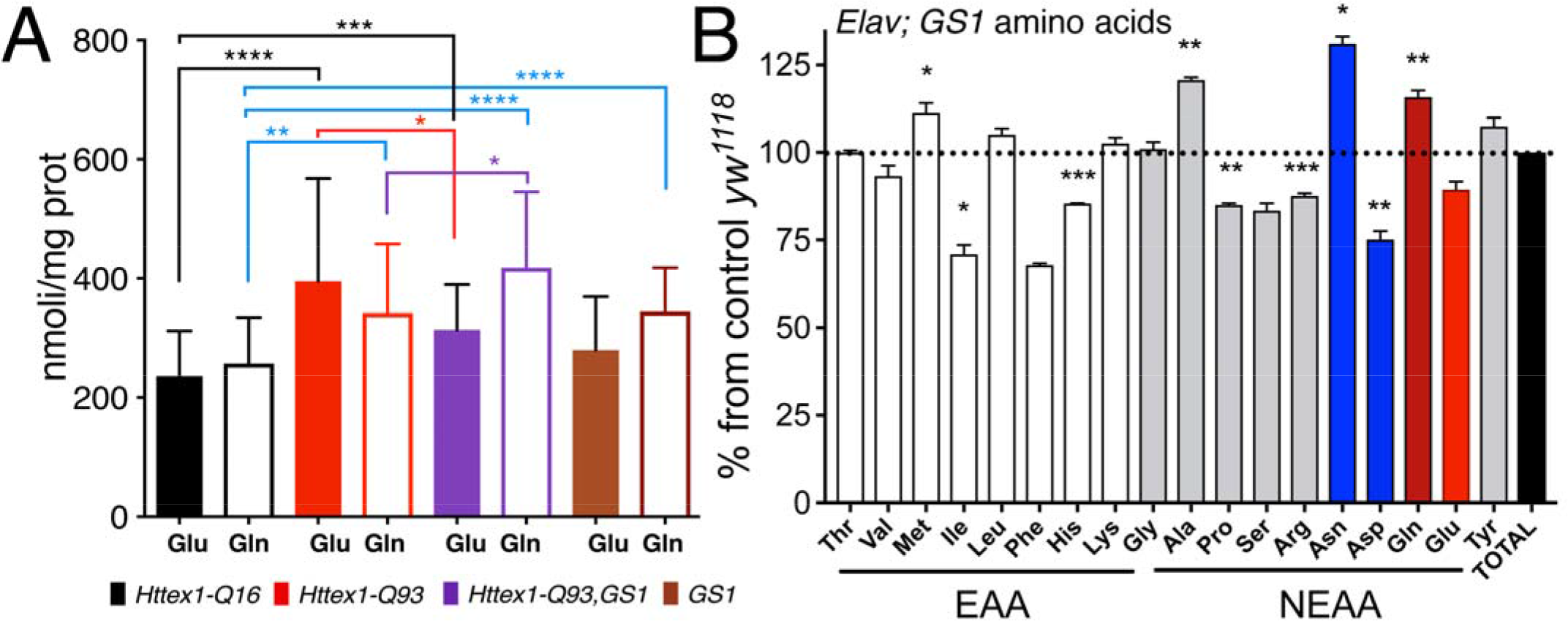
Expression of GS1 in neurons increases Glutamine levels. (A) Quantification of glutamate (Glu) and glutamine (Gln) concentration (MS/HPLC) in lysates from the heads of animals of the indicated genotypes. (B) Amino acids profiling form the heads of animals expressing GS1 in neurons, data are indicated as the % of the concentrations in *GS1* over control *w*^*[1118]*^ animals. EEA= essential amino acids, NEA= non-essential amino acids. The * = *p* < 0.05, ** = *p* < 0.01, **** = *p* < 0.0001 values in panels B were calculated from Student’s *t*-test from two independent experiments, error bars indicate the standard deviations.

To identify significant changes in the pattern of amino acid levels modulated by GS1 expression, we analyzed the profile of Essential (EAA) and Non Essential (NEAA) amino acids in the lysates from heads of flies expressing *GS1* in neurons and compared it to that from control *Httex1-Q16* (Fig. 6B). This analysis confirmed that glutamine (Gln), and alanine (Ala) and methionine (Met) as well, was significantly increased upon GS1 expression (*p*<0.001), while EAA such as isoleucine (Ile) and histidine (His), and NEAA such as proline (Pro) and arginine (Arg) were significantly reduced. Of particular interest was the decrease in the concentration of aspartic acid (Asp) that was accompanied by an increase of asparagine (Asn). This last set of data may suggest the activation in neurons of asparagine synthetase (ASNS), the enzyme that hydrolyzes glutamine to produce asparagine from aspartic acid and glutamate. ASNS is usually elevated in tumor cells to induce cell survival upon nutrient deprivation by switching to the endogenous production of asparagine [27].

### Fibroblasts from patients with Huntington’s disease have decreased autophagy and reduced level of GS1

Since autophagy was found to be defective in HD [37, 38], we analyzed fibroblasts derived from patients expressing the mutant *huntingtin* gene to establish whether the autophagic flux was compromised also in these cells. The amount of cleaved LC3, the human homolog of Atg8a, was determined by immunoblot in fibroblasts from HD patients and control donors. This analysis showed that fibroblasts from HD patients exhibit a reduction of the fast migrating band corresponding to the cleaved LC3-II form as compared to that in control fibroblasts (Fig. 7A, upper gel). A similar result was obtained using anti-Atg8a antibodies (Fig. 7A, middle gel). Since GS1 expression was decreased in *post-mortem* brains from HD patients, we also analyzed its expression and found that GS1 protein is reduced by 43 % in HD fibroblasts (Fig. 7B). Huntingtin is present in both these lines as a band of about 350 kDa; in addition, a band of about 80kDa is detected only in the lysates from HD patients carrying 78-90 CAG repeats in the *HTT* gene (Fig. 7C). Taken together, these results suggest that in fibroblasts carrying mutant *huntingtin* the autophagic flux is compromised and the levels of GS1 are partially reduced. This observation fully supports our observation that GS1 plays an important role in controlling the autophagic flux in *wild-type* cells, but also that its expression may be compromised in pathological conditions, such as HD, which affect the physiological role of autophagy necessary to keep cells, and in particular neurons, in healthy conditions.

**Fig 7:**
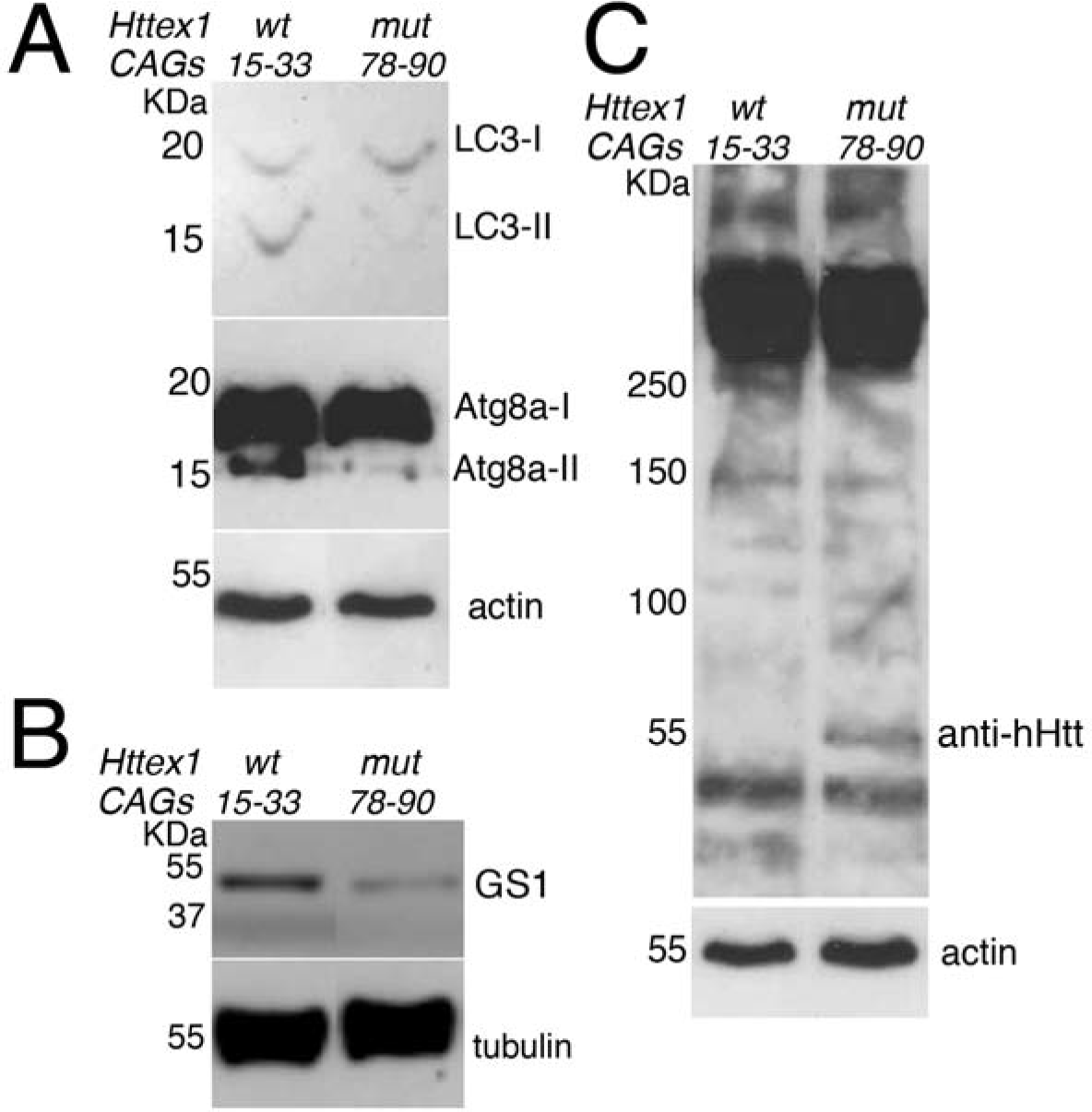
Fibroblasts from HD patients have reduced autophagy and GS1 expression. (A) Western blot showing the cleavage of LC3 and Atg8a proteins in lysates of fibroblasts from control donors with 15-33 CAG repeats and from HD patients with 78-90 CAG repeats (HD). The fast-migrating forms of LC3-II (upper panel) and Atg8a-II (middle panel) are significantly reduced in the extracts from fibroblasts of HD patient as compared to lysates from control donors. Actin is shown as a control for loading (lower panel). (B) Expression of GS1 is reduced in fibroblasts from HD patients; Tubulin was used as a control for loading. (C) Western blot showing human Htt protein expression in fibroblasts from control donors and from patients with 78-90 CAG repeats, Actin was used as control for loading. The antibodies evidence the presence of a band of about 60 kDa that is present only in lysates from cells deriving from HD patient. Experiments were repeated at least twice, data were analyzed from three different patients and a representative situation is shown.

## Discussion

The glutamate-glutamine cycle (GGC) is a sequence of biochemical events between neurons and glia cells necessary for the control of glutamate homeostasis. Previous studies showed that this cycle is impaired in mouse and *Drosophila* models for HD [39, 40]. In addition, the activity of Glutamine Synthetase-1 (GS1), a key regulator of the GGC, was found to be reduced in *post mortem* brains of HD patients [14, 41]. To elucidate the role of GS1 in HD, we modulated its activity in neurons using a *Drosophila* model expressing the mutated human exon1 *Httex1-Q93* that mimics many aspects of neurodegeneration induced by toxic polyglutamine expansion in HD patients [28]. Our data show that neuronal expression of the *Drosophila* homolog of human GS1 ameliorates animal motility and reduces neuronal death induced by expression of *Httex1-Q93*. Moreover, we found that co-expressing GS1 with *Httex1-Q93* reduces the size of the toxic HttQ93 aggregates and increases the levels of macro-autophagy, herein referred to as autophagy, establishing a novel link between the GGC and functional autophagy in post-mitotic cells.

Autophagy is an evolutionarily conserved mechanism that mediates the degradation of cellular debris through the action of lysosomes and has an established positive role in several processes beneficial for neurons under both normal and pathological conditions [1, 42]. Autophagy is activated by the action of specific *autophagy-related genes* (*ATG*s) genes, whose function is highly conserved both in flies and vertebrates [43]. Atg1, the *Drosophila* homolog of mammalian ULK1 kinase, forms the first initiation complex (consisting of the kinase Atg1/ULK1, Atg13, and Atg17) and its activity is inhibited by phosphorylation of TOR kinase [44]. As the process continues, Atg5, which operates downstream in the cascade, becomes involved together with Atg8 in vesicle elongation during the autophagosome maturation. This step is important as lipidation of Atg8 and its proteolytic cleavage are used as readouts of an active autophagic flux in western blots. The autophagosome eventually matures into a closed cargo-containing vesicle, which then fuses with the lysosome to become the autolysosome, where the cargo is degraded for recycling. The latter step is carried out by the protein p62/SQSTM1 or sequestosome-1 that is then degraded by ubiquitination [45]. Recently, a physiological role for the *wild-type* Htt protein was assigned in controlling the initial steps of the autophagosome formation and the recognition of p62 in the cargo formation [46, 47]. Furthermore, the proteolytic stress induced by mutant Htt (mHtt) was shown to favor the interaction of p62/SQSTM1 with ULK1 to regulate the clearance of toxic aggregates [48]. However, when mHtt accumulates over a threshold, the cargo can no longer be eliminated p62/SQSTM1 accumulates blocking the autophagic flux [49]. In our experiments we found that the expression of GS1 in neurons favors the cleavage of Atg8 and reduces the level of p62 also when Httex1-Q93 is co-expressed, suggesting that its activity favors cargo degradation and the autophagic flux.

The important role of autophagy in neurons is revealed by reducing *ATGs* genes, since conditional knock-out of *ATG5* and *ATG7* in neurons results in progressive motoneuron defects, alteration in autophagosome formation and accumulation of ubiquitin-positive protein aggregates in degenerating neurons [50–52]. These phenotypes are conserved in *Drosophila* where *atg7*^*D77*^ mutants have a short life-span, reduced motility, and accumulation of ubiquitinated proteins [53]. Lack of *ULK1*/*ATG1* in mice causes defects in the endocytosis process and abnormal axonal branching, a function conserved also in *C*. *elegans* and *Drosophila* [54, 55]. Moreover, polymorphisms in the *ATG* genes, including *ATG5*, and *ATG7*, have been associated with early onset of NDDs, including HD [56], evidencing the protective role of physiological autophagy also in humans. In our experiments, we found that reducing the expression of *atg1* and *atg5* in neurons of animals expressing *Httex1-Q93* significantly exacerbates the motor degeneration phenotype induced by mHtt, while the rescue driven by GS1 overexpression was completely abolished, further confirming that GS1 function requires a functional autophagic pathway to exert its activity in neurons.

Autophagy is inhibited by nutrients and amino acids that are normally required for TOR activation and translocation to the lysosomal membrane [7, 57]. Our data show that expression of GS1 impairs TOR localization in the lysosome. Such data are supported by the reduction of S6K phosphorylation that was clearly visible in lysates from the heads of animals both expressing *GS1* alone and co-expressing *GS1* and *Httex1-Q93*. GS1 converts glutamate into glutamine, and analysis of the levels of these two amino acids in the heads of animals expressing GS1, showed that glutamate was reduced in *Httex1-Q93*, *GS1* animals where we rather observed a significant increase in glutamine. TOR signaling controls cell growth and protein synthesis in response to nutrients and amino acids. However, its activation depends on the cellular milieu and functions through a mechanism that is mostly conserved from yeast to mammals [58]. In general, amino acid stimulation of TORC1 at the lysosome membrane depends on the activation of the Rag guanosine triphosphatases (GTPases), the Ragulator complex, and the vacuolar H^+^ adenosine triphosphatase (v-ATPase) [59]. However, the ability of single amino acids to control TOR activity seems to be controlled specifically by certain branches of the pathway [60]. Recently, many studies on the ability of different amino acids to stimulate TOR signaling have been focused on the specificity of glutamine [61, 62], leucine [20, 36, 61], and arginine [63].

For example, glutamine, which is a nonessential amino acid synthesized by glutamine synthetase, has been shown to bind or activate the adenosine diphosphate ribosylation factor–1 GTPase (Arf1) to directly activate TORC1 at the lysosome membrane, and in epithelial cells, glutamine requires the function of the v-ATPase to fully activate TORC1 [60]. In addition, glutamine can be used indirectly to stimulate TORC1 via glutaminolysis, where glutamine is first converted into glutamate by GLS (glutaminase). GDH (glutamate dehydrogenase) then converts glutamate into α- ketoglutarate to directly activate TORC1 [64] or to participate in the anaplerotic reactions of the tricarboxylic acid cycle when nutrients are reduced [65, 66].

Because of its involvement in a multiplicity of metabolic pathways, the function of glutamine in the activation of TOR signaling still presents some controversial results. Glutamine has been shown mainly to activate TORC1 directly or by a positive feed-back loop controlled by autophagy that increases the pool of amino acids, including glutamine and arginine to re-activate the TOR pathway [67]. Our data in neurons show that TOR signaling is reduced by expression of GS1 (low level of S6K phosphorylation, and reduced TOR localization in the lysosome), even when glutamine levels increase, suggesting that in terminally differentiated cells GS1 may induce a condition of “starvation-like” that favors autophagy without reactivation of TORC1 even in the presence of glutamine. In support of this mechanism, van der Vos and colleagues showed the presence of a similar mechanism in cancer cells under nutrient starvation condition, where the transcriptional induction of GS1 by FOXO3 resulted in autophagy and cell survival with the concomitant inhibition of the TOR pathway [68].

Glutamine is utilized by the cells also to control the import of essential amino acids (EAAs). Indeed, a distinct mechanism described so far in epithelial cells, links leucine and EAAs to the glutamine flux [61]. Through this mechanism, glutamine imported in the cytoplasm by the SLC1A5 transporter is exported by the heterodimeric anti-porter system (SLC7A5-SLC3A2) to import leucine and ultimately stimulate TORC1 [69]. Moreover, since our amino acid analysis from the heads of animals expressing GS1 in neurons shows an increase in glutamine levels we can speculate that the system that controls the export of glutamine is impaired, and consequently this may reduce the import of EAAs such as isoleucine (Ile) and histidine (His) resulting in the inhibition of TOR signaling.

Expression of GS1 also reduces the level of glutamate in favor of glutamine. Glutamate is a non-essential amino acid that function as an excitatory neurotransmitter in the central nervous system and as a substrate in many distinct reactions [70]. Glutamate is synthesized from glutamine, α-ketoglutarate and 5-oxoproline and it serves as a precursor molecule for the biosynthesis of various amino acids including proline (Pro) and arginine (Arg) [65, 66]. Interestingly, in the heads of animals expressing GS1, we found a significant reduction of Pro and Arg, both relevant in the mechanism of TOR activation through specific amino acid sensors [59]. In particular, the lysosomal amino acid transporter SLC38A9, proposed in mammals to function upstream of TORC1 [26], was found necessary for TOR activation by Arg, while its constitutive expression renders the cells insensitive to the stimulation by amino acids, including arginine [63]. So far there are no data describing the mechanism of activation by SLC38A9 in neurons. However, since we found that Arg levels are reduced in the heads of *elav*>*GS1* animals, we hypothesize that in neurons Arg is required for the activation of TOR by SLC38A9. Moreover, these data may also suggest that in neurons activation of SLC38A9 by Arg acts downstream of glutamine in the activation of the TOR pathway since glutamine is not a limit. It is worthy to note that, although the *Drosophila* SLC38A9 transporter has not been characterized yet, the *Drosophila* genome contains three genes expressed in the brain (CG13743, CG30394, and CG320181) [71–73], whose sequences are significantly homologous to that of human SLC38A9 [74], suggesting that they may encode the fly functional homolog(s) of SLC38A9.

Another mechanism that cells use to counteract the decrease of glutamate levels, is to convert the excess of glutamine into asparagine through the enzyme asparagine synthetase (ASNS), which converts aspartate (Asp) and glutamine to asparagine (Asn) and glutamate (Glu). By looking at the concentration of those amino acids in the heads of animals expressing GS1, we also found that Asn concentration was significantly increased while, on the contrary, Asp decreased. This would indicate an upregulation of the activity of ASNS, an enzyme that is induced in tumor cells during nutrient starvation [26, 27, 75]. This may suggest that ASNS in neurons responds to amino acid starvation induced by GS1 by increasing the levels of Asn and Glu, which is also in the substrate of GS1. In support to this hypothesis, we found that upon GS1 expression Glu levels are only slightly reduced, while the concentration of few other amino acids, such as isoleucine, phenylalanine, histidine, proline, and serine, that are necessary to TOR activation are significantly lower. Alanine levels instead increased, possibly because of the rise in glutamine that indirectly participates in glutamate synthesis via glutaminolysis, or to support metabolic pathways in the biosynthesis of proteins [76].

## Conclusion

Our data show a novel function for GS1 in controlling autophagy in neurons to ameliorate the detrimental effect of the accumulation of toxic Htt aggregates. Neurons differ from other cell types for being post-mitotic and highly dependent on the endo-lysosomal pathway for active signaling in axons and dendrites. Therefore, these cells particularly require effective protein degradation as a quality control mechanism for cell survival. Any alteration of this process, especially under disease conditions, causes the accumulation of abnormal proteins, leading to cellular toxicity and ultimately neurodegeneration. Malfunctions in components of the glutamate-glutamine cycle have been detected in many neurodegenerative diseases including Parkinson, Alzheimer and Huntington [14, 77, 78]. Significant progress has been made to understand the nature of both ageing and neurodegeneration, and the impairment of autophagy is one of the major risk and possible cause of numerous NDDs that are diagnosed in elderly when the physiological role of autophagy starts to decline. Here, we were able to find a novel function for GS1, a key enzyme that controls glutamate homeostasis, in the regulation of autophagy in neuron [79]. Moreover, our observations in fibroblasts from HD patients suggest that the link between GS1 activity and autophagy is conserved in humans as well, thereby paving the way for future research on the role of GS1 in autophagy and in potential HD therapeutic targets using HD *Drosophila* models.

## Materials and Methods

### Fly husbandry and lines

Animals were raised at low density, at 25°C, on a standard food medium containing 9 g/L agar (ZN5 B&V, Italy), 75 g/L corn flour, 60 g/L white sugar, 30 g/L brewer yeast (Acros Organic), 50g/L fresh yeast, and 50 ml/L molasses (Biosigma), along with nipagin and propionic acid (Acros Organic). The following fly lines were obtained from the Bloomington *Drosophila* Stock Center: *UAS-Httex1-16Q* (B33810), *{EP}-GS1* is an enhancer-promoter (EP)-line with a random insertion of the *P*-element, carrying an *UAS* enhancer sequence and a basal promoter, in the *GS1* locus (B27940), *UAS-GS1RNAi* (B40836), *UAS-mCherry-Atg8a* (B37750), *elav-Gal4*^*C155*^ on the *X ch*. (B458) and *GMR-Gal4* (B9146). Fly lines bearing the: *UAS-Httex1-Q93* was a gift of from L. Marsh (University of California, Irvine), *UAS-LAMP1-GFP* and *UAS-Atg1* were a gift from F. Parisi (Beatson Institute for Cancer Research, Glasgow) and *UAS-Atg5-RNAi* was a gift from T. Neufeld (University of Minnesota).

### Survival at eclosure analysis

The number of animals and their timing of eclosion from every cross with the *elav-Gal4*^*C155*^ driver were quantified and the percentage of animals eclosed was calculated for each of the genotypes of interest with respect to the expected number of animals that eclosed.

### Motility assays

The motility assay with larvae was performed with ten 3^rd^ instar animals for each genotype of interest. Larvae were collected, washed with PBS and then transferred to a 14 cm Petri dish containing 1% agarose in PBS where a visible grid was drawn (Supplementary Fig. 3). The number of lines crossed by one larva in 1 minute at room temperature was counted. Statistical analysis was performed with the Student’s *t*-test; values for each genotype are represented as mean ± standard deviation (SD). Experiments were repeated at least three times. For motility assays with adult animals, one day-old females of each genotype were transferred in a plastic vial without food (15 animals for each genotype), and their ability to climb up the empty vial after a knock-down to the bottom was analyzed, as previously described [80]. The number of flies that were able to climb half of the tube in 15 seconds was recorded. Values were expressed as percentage of success with respect to the total number of flies in the vial. For each genotype the test was repeated 30 times for each timepoint. After the test, adults were transferred in vials with food and vials were changed every two days. Data are represented as a curve of progressive motility impairment. The statistical analysis of variance (one-way ANOVA) was performed using PRISM GraphPad Software (CA). Every experiment was repeated at least 6 times.

### Quantification of fluorescence (GFP) on the adult compound eye

Photographs of adult eyes expressing the indicated UAS-transgenes in the retina and co-expressing *UAS-GFP* using the *GMR-Gal4* promoter, were taken at 8 days after eclosion using a Leica stereomicroscope MZ10F with a fluorescent source and a camera, using 5x magnification. The intensity of fluorescence in each image was quantified as integrated density from an area containing 20 ommatidia using Adobe Photoshop-CS4 and expressed as GFP intensity/area,. At least 10 eyes (one for each animal) were used for each genotype and experiments were repeated twice.

### Western blot

Proteins were extracted from ten brains of larvae or 10 adult heads collected in lysis buffer (50 mM Hepes/NaOH pH 7.4, 250 mM NaCl, 1 mM EDTA, 1.5% Triton X-100 containing phosphatases and proteases inhibitors (Roche). For detection of the Huntingtin protein, lysates were extracted using 2% SDS lysis buffer (50 mM Hepes/NaOH pH 7.4, 250 mM NaCl, 1 mM EDTA, 2% SDS containing phosphatases and proteases inhibitors. Samples were sonicated three times for 10 seconds using a Branson Ultrasonic Sonifier 250 (Branson, Darbury, CA, USA) equipped with a microtip set at 25% power. Tissue and cell debries were removed by centrifugation at 10000 g for 30 min at 4°C. Proteins in the crude extract were quantified by BCA Protein assay Reagent Kit (Pierce), following the manufacturer’s instructions with bovine serum albumin as the standard protein. For SDS-PAGE, samples were incubated for 8 min at 100°C in standard reducing loading buffer; 40 μg of total protein were run on a SDS-polyacrylamide gel and transferred onto nitrocellulose membranes (GE Healthcare). After blocking in 5% (w/v) non-fat milk in TBS-0,05% Tween (TBS-T), membranes were incubated overnight with primary antibodies against hHTT (1:500, Abcam ab109115), Elav (1:100, Hybridoma Bank #9F8A9), Synapsin (1:100, Hybridoma Bank #8C3), Actin5c (1:200 Hybridoma Bank #JL20), Tubulin (1:500, Chemicon, MAB3408), GABARAB/Atg8a (1:500 ABCAM ab109364), mTOR (1:500 Abcam ab45989), anti TOR-P1(1:400 gift from A. Teleman, DKFZ, Heidelberg), GS1 (1:800 Abcam ab64613), SQSTM1/p62 (Cell Signaling 5114), anti Drosophila p70 S6K (Cell Signaling 9209), ULK1/ATG1 (1:500 Cell Signaling #D8H5), or ATG5 (1:500 Thermo Fisher #OSA00026W).

### Filter trap assay

Ten larval brains or adult heads for each genotype were homogenized in lysis buffer (50 mM Tris-HCl pH 6.8, 2% SDS plus proteases and phosphatases inhibitors). Fifty micrograms of protein extracts were loaded on a dot-blot device and filtered through cellulose acetate membrane (0.22 μm pore size), previously washed in 1% (w/v) SDS solution in PBS. After filtration, membranes were washed three times in TBS-T, saturated in 5% non-fat milk and processed for immunodetection with anti Htt antibodies from Abcam, as described above.

### Patient-derived cell lines

Each individual providing a biological sample signed written informed consent approved by the Institutional Review Board of the Fondazione IRCCS Istituto Neurologico Carlo Besta, Milan, Italy, in agreement with the Declaration of Helsinki. Fibroblasts from patient and controls were grown in the DMEM-high glucose or -galactose media as described previously in [86].

### MS/HPLC for glutamate and glutamine quantification and analysis of amino acids

10 heads from animals at 8 DAE were resuspended in 100 μl of Hepes/NaOH 50mM with proteases and phosphatases inhibitors (Roche), samples were disrupted using a mechanical tip following two rounds of sonication of 10 sec each. After centrifugation at 13000 rpm for 30 min at 4°C, 20 μl for each sample were set aside for Bradford analysis, while supernatants were diluted in buffer containing DL-Lysine D_4_ and DL-Ornitine D_6_ as internal standards (Cambridge Isotopes Labs, Inc Eurisotop) and 2/3 v/v of methanol, to precipitate the proteins. After centrifugation aminoacid analysis was performed on a Perkin Elmer Series 200 HPLC equipped with 4 *μ*m, 2.0 × 75 mm Synergi Polar-RP (Phenomenex) column and coupled with triple quadrupole API2000 tandem mass spectrometer (AB SCIEX, MA, USA). AB SCIEX Analyst software (version 1.4) was used for data acquisition and analysis. Analysis of free amino acids was performed by Innovhub, Stazioni Sperimentali per Industria, Milan (Italy) with the following protocol: 30 heads of each genotype were resuspended in 300 μl of HPLC-grade H_2_O, samples were disrupted using a mechanical tip following two rounds of sonication of 10 sec each. 20 μl were set aside for Bradford analysis, while 75 μl of a 6% solution of sulfosalicylic acid were added to the final concentration of 1.5%. Samples were centrifuged for 20 minutes at 13.0000 rpm and supernatants were used for the analysis. The protocol applied (UNI 22615:1992) uses ionic exchange chromatography where the amino acids are separated by chromatography using an automatic Lithium High Performance Physiological Column with the instrument Biochrom 30^+^ from Biochrom. The separation is based on using Lithium Citrated buffer solutions using a five steps pH separation from pH 2.80 up to pH 3.55. Amino acids were visualized on TLC using ninidrin and detected at 440 e 570 nm.

### Quantification of GS1 activity

*Drosophila* larvae or heads that had been flash frozen in liquid nitrogen were resuspended in 200 μl 50 mM Hepes/NaOH buffer, pH 7.4 and 0.5% protease inhibitors cocktail (Sigma P8849). They were homogenized with a Branson 250 sonifier equipped with a microtip (25% power, 20% duty cycle) by applying four series of 10 sonication pulses and controlling the temperature by immersion in an ice bath. The homogenate was centrifuged in a microfuge at 13000 rpm for 30 min at 4°C and passed through a Ultrafree-MC (Millipore UFC306V) filter device by centrifuging at 11000 rpm for 15 min at 4°C. The filtrate was used for protein and activity assays. Protein concentration was determined with the Bradford method and bovine serum albumin as the reference protein [81].GS activity was assayed by quantifying the amount γ-glutamylhydroxyamate (GGHA) produced as described in [82–84]. In preliminary experiments a range of GGHA concentrations (0.07-0.15 mM) was used to determine the sensitivity and linear range of the assay. Briefly, a known amount of crude extract (0.1-0.5 mg in up to 120 μl) was incubated for 1 h at 37°C in 50 mM Tris/HCl buffer, pH 7.5, hydroxylamine (40 mM final concentration), MgCl_2_ (40 mM) L-glutamate (20 mM) and ATP (5 mM) in a final volume of 600 μl. The reaction was blocked by adding 200 μl of a FeCl_3_ solution (3.3% FeCl_3_, 8% trichloroacetic acid (TCA), 0.67 M HCl) and centrifuged at top speed for 10 min. The amount of GGHA produced was calculated from the absorbance value at 500 nm and 600 nm with a HP 8453 diode array spectrophotometer using the reported extinction coefficients of 1.004 mM^−1^cm^−1^ and 0.471 mM^−1^ cm^−1^[82] for the crude extracts derived from larvae and heads, respectively; by monitoring the entire spectrum of the final solution, we ensured that no artifacts arose from turbidity, precipitation or presence of pigments, especially with *Drosophila* heads. For each assay a blank devoid of L-Glu and ATP was set up and the measured absorbance was subtracted to that of the corresponding sample prior to calculation of the activity value. The amount of larvae or heads extracts were selected in order to ensure that GGHA concentration increase was linear with respect to the incubation time. L-Methionine sulfoxide (MSO) was added to reaction mixtures at a final concentration of 1 mM (from 10 mM stock solution in 50 mM Hepes/NaOH buffer, pH 7.4) to test its effect on GS1 activity. Activity was expressed as milliunits/mg protein, where 1 unit of GS activity is the amount of GS that converts 1 μmol L-Glu into product per minute. GS activity detected in wild-type larvae was taken as 100% activity to normalize the values obtained with different larvae batches and in comparison with the values measured with transfected larvae. For each sample, replicates differed by less than 15%.

### Immunofluorescences and quantification of the size of Htt-Q93 aggregates

To visualize human mutant Htt-Q93 aggregates, females *elav-Gal4; UAS-GFP* females were crossed with males carrying the appropriate UAS-transgenes. 3^rd^ instar larvae were collected and washed in PBS. Brains were dissected and fixed for 30 minutes in 4% paraformaldehyde (PFA) on ice. After tissues permeabilization with 0.5 % Triton X-100, samples were washed in PBS-0.1% Tween20 (PBST) and blocked in 3% BSA (in PBST) for 30 minutes on ice. Samples were incubated with anti-Htt antibody in 3% BSA overnight at 4°C. After 3 washes of 10 minutes each in PBST samples were incubated with a mouse or rabbit secondary antibodies Alexa 555 (Invitrogen) 1:400 in 3% BSA for 2 hours at RT on the shaker. After extensive washing with PBST, brains were mounted on slides in 20 μl of Mowiol. Nuclei were stained with Hoechst 33258 added at the final concentration of 1μg/ml. The integrated density of the fluorescence was acquired from at least 10 confocal’s *z*-stacks in the specific brain area and quantification of the integrated density was considered directly proportional to the aggregate size. Aggregate density was quantified by analyzing the integrated density of the signal from the confocal stacks using *ImageJ* 1.50i software (Wayne Rasband NIH, USA).

### Co-localization analysis

To analyze TOR co-localization with lysosome, females *elav-Gal4; UAS-LAMP1-GFP* females were crossed with males carrying the appropriate UAS-transgenes. 3^rd^ larvae were collected, washed and brains were dissected and processed as described above, incubated with anti mTOR antibodies 1:200 overnight at 4°C and processed as above using anti-rabbit secondary antibody Alexa 555 (Invitrogen) at 1:400 in 3% BSA, and then processed as described above. Images were acquired at the confocal microscope (SM5 LEICA Leserteknik GmbH) at different magnifications. For each genotype, the acquisition of confocal images were from at least 4 separate brains and the statistics was calculated on 10 *z*-stacks images/brain. Co-localization of TOR and LAMP1-GFP was quantified using ImageJ 1.50i software using JACoP plugin, Pearson’s algorithm with Manders Coefficients and Costes automatic threshold (MCC) [85].

### Statistical analysis

Analysis of the variance was calculated using one-way ANOVA with Tukey multi-comparisons test. Student *t*-test analysis between genotypes was performed using GraphPad-PRISM7. One to four asterisk indicate: *p*\* < 0.05, *p* **< 0.01, *p*\*\*\* < 0.001, *p* ****< 0.0001, respectively.

### Autophagy analysis

*elav-Gal4; UAS-GFP; UAS-mCherry-Atg8a* females were crossed with males carrying the appropriate UAS-transgenes. Third instar larvae (72-80 hours after hatching) were collected, washed, and brains dissected in phosphate-buffered saline (PBS) buffer. Brains were fixed for 30 minutes in 4% paraformaldehyde (PFA) on ice. Nuclei were stained with Hoechst 33258, samples mounted in Mowiol and images acquired using a confocal microscope (SM5 LEICA Leserteknik GmbH) using *ImageJ* 1.50i software or *AdobePhotoshop-CS4*. The quantification of autophagy in the *mCherry-Atg8a* vesicles, representing autophagosomes, was analyzed by measuring the intensity of fluorescence inside a fixed area in al least four different regions from each brain, and in four animals for each genotype.

## Acknowledgements

We thanks Alessia Soldano (University of Trento) and Chiara Zuccato (University of Milan) for reading the manuscript. The Confocal Facility at Istituto FIRC per Oncologia Molecolare (IFOM) Milan, (Italy). Sara Leo, Michela Roccuzzo and Giorgina Scarduelli at the Advanced Imaging Core Facility at Department-CIBIO University of Trento, for helping with the confocal acquisition and statistical analysis. Aurelio Teleman, Deutsches Krebsforschungszentrum (DKFZ) Heidelberg (Germany) for the anti *Drosophila* TOR antibody.

## Author contributions

LV, CP, GL, TV, SS, MR, VM, MR, MRG, MEP, MAV, PB performed the experiments. LV, CP, MEP, DG, MAV FT, PB wrote the manuscript MEP, MAV, CG, FT and PB designed experiments.

