## supplemental figures for "Glutamine Synthetase-1 induces autophagy-lysosomal degradation of huntingtin aggregates and ameliorates animal motility in a *Drosophila* model for Huntington’s disease"

### Supplementary figures:

#### Supplementary Figure 1A:

*ClustalW* amino-acid sequence alignment of *Drosophila* (UniProt E1JHQ1) and human (UniProt P15104). The two *GS1* genes show 70.5 % of identity on 298 aminoacids. The common ligand binding residues are indicated with purple (ATP), blue (glutamate) green (ammonia), yellow (metal coordination) [1].

```
Hu       -------------------------------------

Dm 1     MALRVAGLFLKKELVAPATQ-QLRLLRTGNTTR-SQF   35

Hu 1     MTTSASSHLNKGIKQVYMSLPQ-GEKVQAMYIWIDGTGEGLRCKTRTLDSEPKCVEELPE   59
            S ++ L+K I Q Y +L     +VQA Y+WIDGTGE +R K R LD  P  VE+LP+
Dm 36    LANSPNTALDKSILQRYRNLETPANRVQATYLWIDGTGENIRLKDRVLDKVPSSVEDLPD   95

Hu 60    WNFDGSSTLQSEGSNSDMYLVPAAMFRDPFRKDPNK-LVLCEVFKYNRRPAETNLRHTCK   118
         W +DGSST Q+ G NSD  L P A++RDPF+   N  +VLC+ +  + +P  +N R   +
Dm 96    WQYDGSSTYQAHGENSDTTLKPRAIYRDPFKPGKNDVIVLCDTYSADGKPTASNKRAAFQ   155

Hu 119   RIMDMVSNQHPWFGMEQEYTLMGTDGHPFGWPSNGFPGPQGPYYCGVGADRAYGRDIVEA   178
+           D++S+Q PWFG+EQEYTL+  DG PFGWP NGFP PQGPYYCGVGADR Y RD+VEA
Dm 156   AAIDLISDQEPWFGIEQEYTLLDVDGRPFGWPENGFPAPQGPYYCGVGADRVYARDLVEA   215

Hu 179   HYRACLYAGVKIAGTNAEVMPAQWEFQIGPCEGISMGDHLWVARFILHRVCEDFGVIATF   238
         H  ACLYAG+  AGTNAEVMPAQWEFQIGP  GI   D LWV+R+IL R+ E++GV+ TF
Dm 216   HVVACLYAGIDFAGTNAEVMPAQWEFQIGPA-GIKACDDLWVSRYILQRIAEEYGVVVTF   274

Hu 239   DPKPIPGNWNGAGCHTNFSTKAMREENGLKYIEEAIEKLSKRHQYHIRAYDPKGGLDNAR   298
         DPKP+ G WNGAG HTNFSTK MR + G+K IEEAIEKLSKRH+ HI+AYDPK G DN R
Dm 275   DPKPMEGQWNGAGAHTNFSTKEMRADGGIKAIEEAIEKLSKRHERHIKAYDPKEGKDNER   334

Hu 299   RLTGFHETSNINDFSAGVANRSASIRIPRTVGQEKKGYFEDRRPSANCDPFSVTEALIRT   358
         RL G  ETS+I+ FS GVANR+ S+R+PR V    KGY EDRRPS+NCDP++V  A++RT
Dm 335   RLVGRLETSSIDKFSWGVANRAVSVRVPRGVATAGKGYLEDRRPSSNCDPYAVCNAIVRT   394

Hu 359   CLLNETGDEPFQYKN   373
         CLLNE
Dm 395   CLLNE----------   399
```

**Supplementary Figure 1B:**

***Analysis of GS1-mRNA* in whole larvae expressing *UAS-GS1* under the *actin-Gal4* promoter**. Quantitative RT-PCR from larvae expressing the *{EP}-GS1*$^{C3347}$ transgene using the *actin-Gal4* promoter shows the indicated transcript level in *actin>EP-GS1* and control animals; *actin5C* was used as the internal control. *$p$<0.05, values were calculated from Student's $t$-test from at least three independent experiments, error bars indicate the standard deviations.

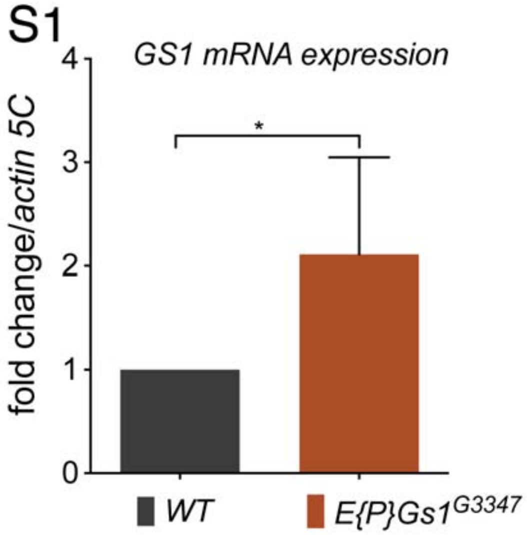

**Supplementary Figure 2:**

**(A) Reduction of GS1, using GS1-RNAi, enhances retinal degeneration induced by the expression of *Httex1-HttQ93*.**

Photographs of eyes of adult female flies at 8 days after eclosion (DAE) expressing (left) *UAS-Httex1-HttQ93* in combination with *UAS-GS1-RNAi* using the retina-specific *GMR-Gal4* promoter, that is expressed in late in 3rd instar in the differentiated ommatidia. These animals present an enhancement of their retinal degeneration that at 8 DAE that was comparable to that of *GMR-UAS-Httex1-Q93* animals at 20 DAE (right).

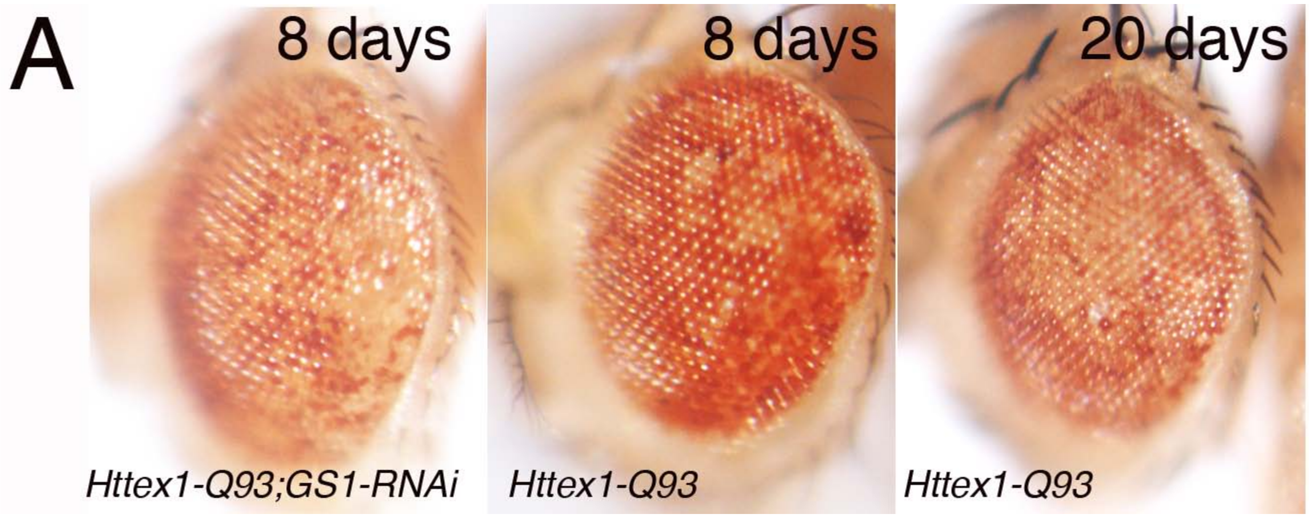


**(B) Retinal degeneration is visualized in the adult eyes using *GMR-GFP*.** Lateral view of eyes of adult females expressing *GMR-GFP* and the relative transgenes, photographed at 8 DAE, showing the relative intensity of GFP.

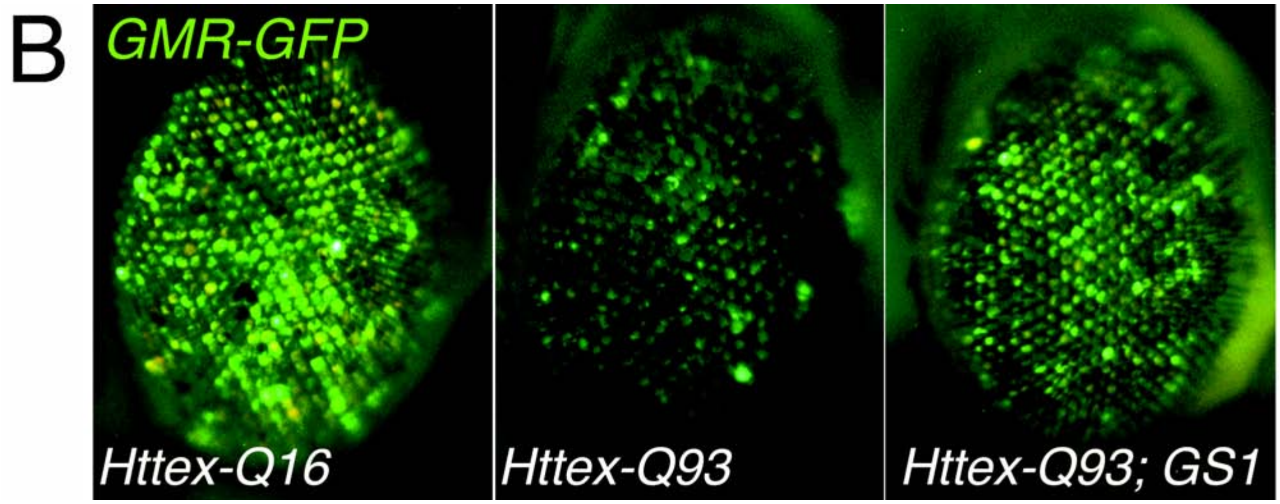

**Supplementary Figure 3: Time line and schematic representation of the motility assays in larvae (MOVIE) and adults (MOVIE) expressing the *elav$^{c155}$-Httex1Q93* and control *elav$^{c155}$-Httex1Q16* transgene**.

(A) Schematic representation of the N-terminus of the Huntingtin protein showing the expanded CAG 16-93 sequence (black). (B) Time line of the appearance of the motility defects caused by the expression of the *Httex1-Q93* gene using the *elav-Gal4* promoter; the motor defects and the presence of Htt-aggregates are already detected after 24-48 hrs after egg laying (AEL)**.** E =embryos, L1, L2 and L3 = first, second and third larval instar respectively. **(C) Petri dish used for the motility assay of larvae.** A grid was drawn on the Petri dish containing 1% agarose in 1X PBS, the red spot is where each larva is put at the beginning of the test. The number of lines crossed by each larva in 1 minute was counted. (D-E) and MOVIE showing the motility of *elav-Httex1-Q16* (E) and *elav-Httex1-Q93 (F)* 3$^{rd}$ instar larvae. (F) Movie showing adult climbing assay of *elav-Httex1-Q16* and *elav-Httex1-Q93* animals at 15 DAE.

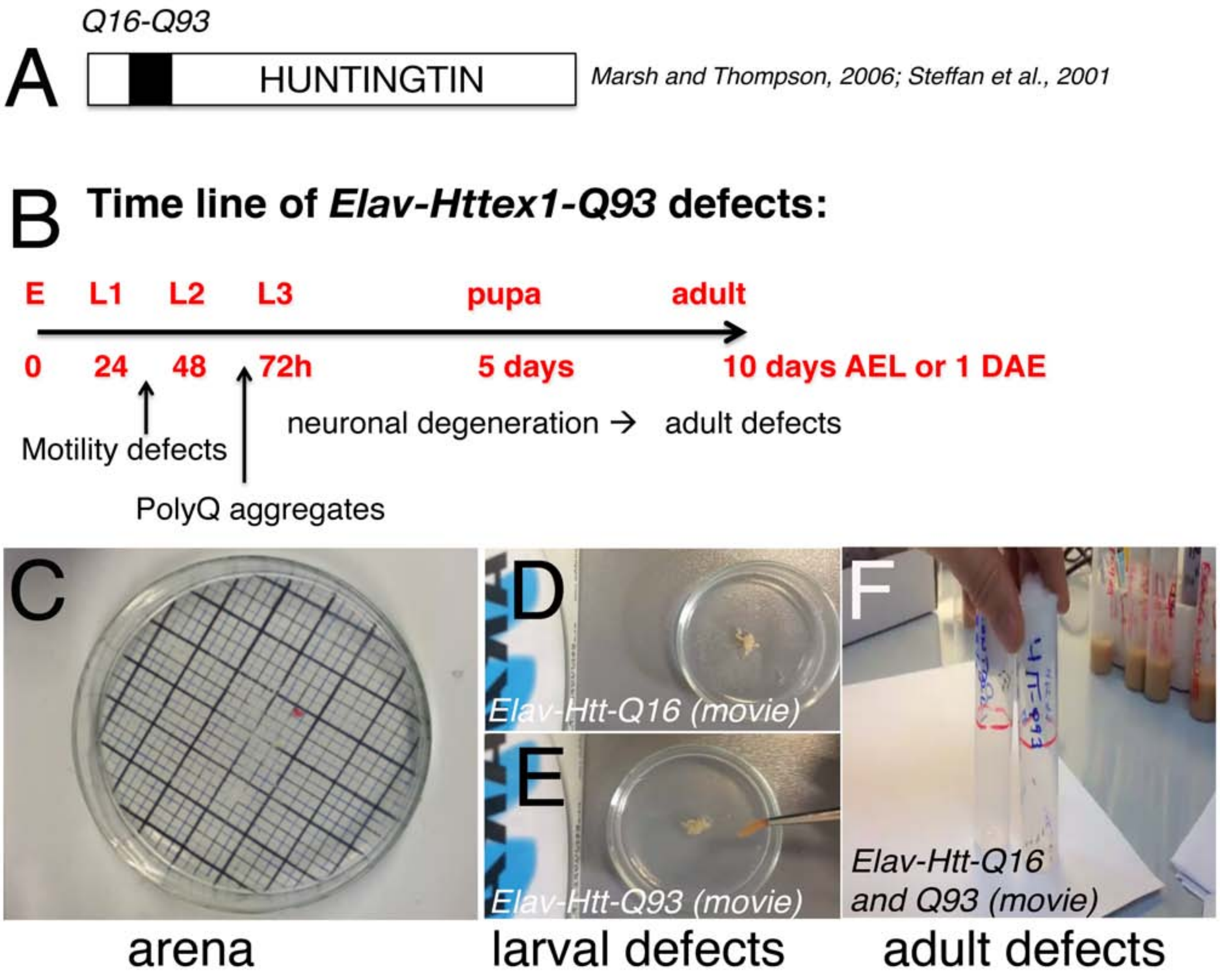

**Supplementary Figure 4: MSO (methionine sulfoximine) specifically inhibits GS1 activity in heads' lysates from from *elav*$^{c155}$ control *and elav*$^{c155}$*-; GS1* females.**

Lysates from heads of animals of the indicated genotype were treated with the specific GS1 inhibitor MSO at 1mM final concentration prior analysis of GS1 enzymatic activity as described in Material and Methods.

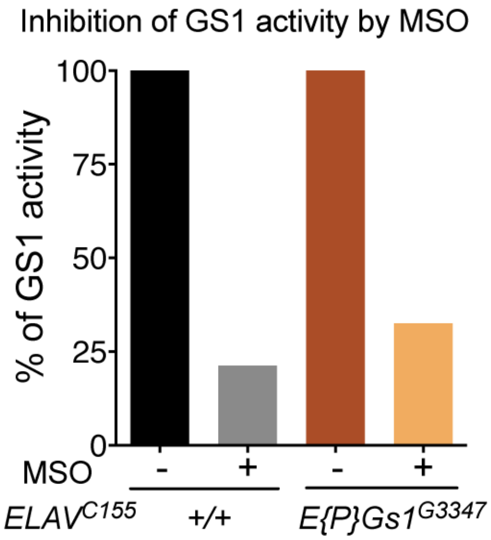

**Supplementary Figure 5:**

**In adult males, co-expression of *GS1* with *Httex1-Q93* in neurons partially rescues animal survival (A) but not climbing defects (B) induced by *elav$^{c155}$*-*Httex1-Q93* in adults males.**

Lethality in adult males was scored over time. Males expressing *Httex1-Q93* (red) showed 50% of lethality at about 5 DAE, while their climbing activity was significantly reduced with 50% reduction at about 2-3 DAE, compared to control *Httex1-Q16* males (black) (ANOVA $P<0.0001$). Co-expression of GS1 was able to partially rescue these defects, as shown in *Httex1-Q93,GS1* (purple) one-way ANOVA analysis between *Httex1-Q93* and *Httex1-Q93,GS1* was significant only for the survival ret showed (ANOVA $P<0.05$).

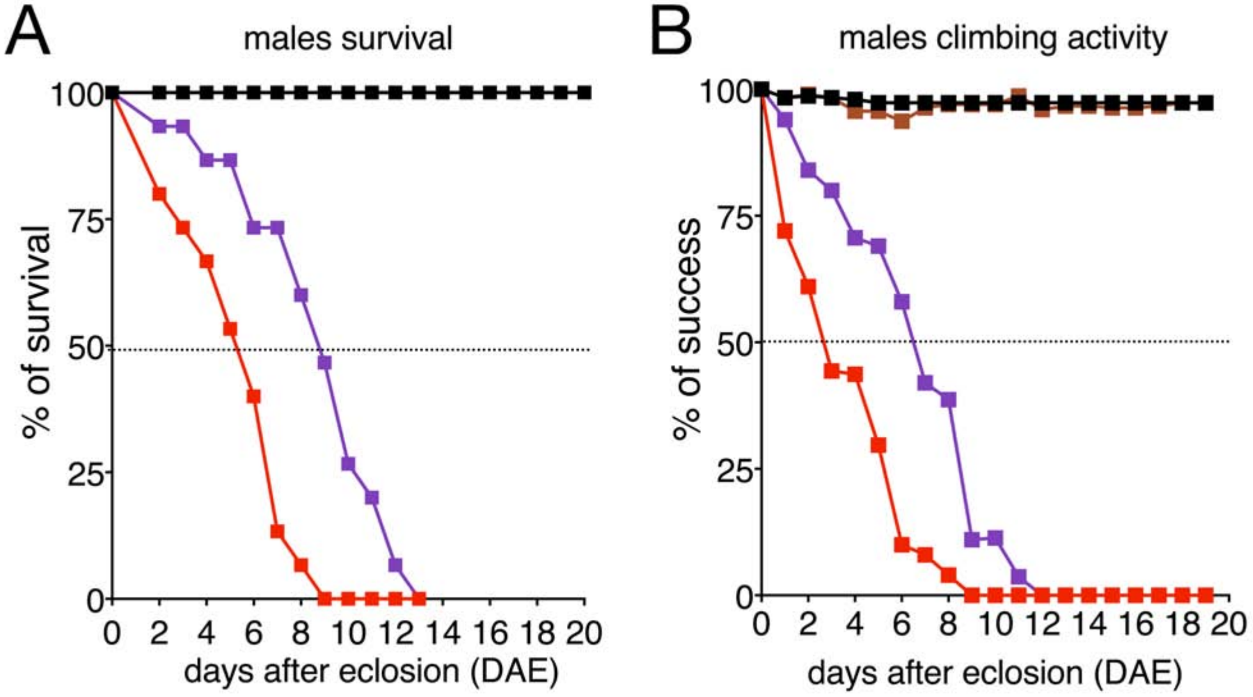

**Supplementary Figure 6:**

**Co-expression of *GS1* with *Httex1-Q93* in neurons partially rescues animal lethality (A) and climbing defects (B) induced by *elav$^{c155}$-Httex1-Q93* in adult males using two others independent recombinant *Httex1-Q93*, *GS1* lines.**

We analyzed defects in climbing activity in adult females, using three independent lines where *GS1* was recombined with *Httex1-Q93*. Expression of *Httex1-Q93* significantly decreased the motility of animals compare to control *Httex1-Q16,* ANOVA: *Httex1-Q16 vs. Httex1-Q93* $P<0.0001$. Expression of *GS1* ameliorate these defects as seen in *Httex1-Q93, GS1 line 1,* ANOVA: *Httex1-Q93* vs. *Htt-Q93, GS1* (1) $P<0.05$); or *Httex1-Q93, GS1 line 3,* ANOVA: *Httex1-Q93* vs. *Htt-Q93, GS1* (3) $P<0.01$), and *Httex1-Q93, GS1 line 4,* ANOVA: *Httex1-Q93* vs. *Htt-Q93, GS1* (4) $P<0.05$). See supplementary Excel file.

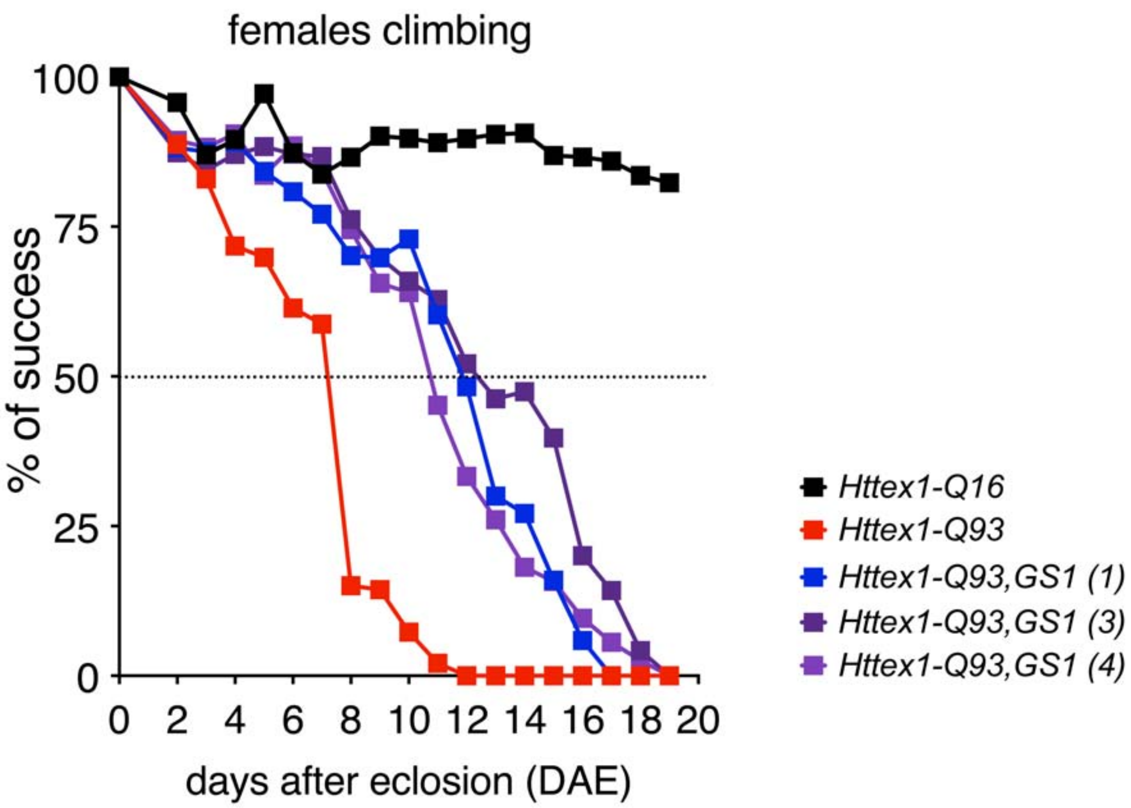

**Supplementary Figure 7: Efficiency of *Atg1-RNAi* and *Atg5-IR 24-1* in reducing relative protein expression in adult heads.** (A-D) Reduction of *Atg1* decreases autophagy in neurons. Quantification from confocal images of brains from 3rd instar larvae expressing *elav-Gal4; mCherryAtg8a* in the neurons of the calyx to mark autophagosomes. (A) Integrated density corresponding to autophagosome density (see Materials and Methods) of mCherry measured in the calyx from: control *elav>w$^{1118}$*(B-black), *elav>mCherryAtg8a-RNAi* on the II chromosome (C-gray), *elav>mCherryAtg8a-RNAi* on the III chromosome (D-dark-gray) animals. Analysis of mCherry shows a significant decrease of Atg8a levels in C and D, if compared to control A. There is no significant difference in Atg8-mCherry levels between samples expressing *Atg1-RNAi* (II) or *Atg1-RNAi (III)*. (One-way ANOVA test, error bars – SEM, ** p< 0,01; *** p< 0,001)

(E) Western blots showing the expression of ATG1 (left) and ATG5 (right) proteins using the promoter *repo-Gal4* in lysates from the heads of females. Repo is a promoter for glia and we use it as a control for the efficiency of *Atg1-Rnai* and *Atg5-RNAi* lines in a set of experiments involving glia. Here we used it here to show the efficiency of our lines. Tubulin was used as loading control.

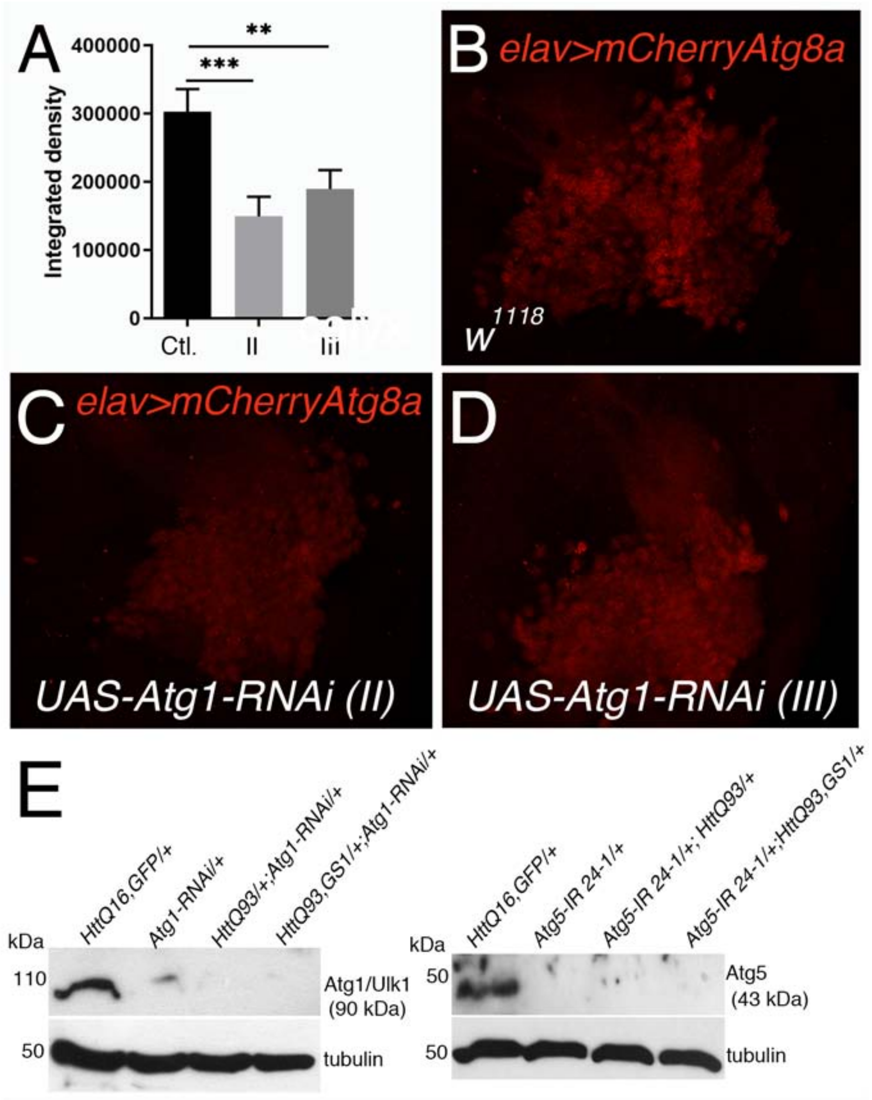

**Supplementary Figure 7: Characterization of anti Tor (target of rapamycin) antibody. (A-F) Immunostaining of third instar imaginal disc expressing *tor-RNAi* using *engrailed-Gal4.*** Photographs of immaginal wing discs from 3rd instar larvae after staining with anti-TOR antibody, where using the *engrailed-Gal4* promoter, Tor expression was reduced in the posterior compartment by expression of the *UAS-tor-RNAi*, membrane-*UAS-GFP* was also co-expressed to mark the cells in the compartment (A). TOR antibody evidenced a clear reduction in TOR protein in the posterior compartment of the disc marked by GFP (A) where *tor-RNAi is* expressed. This difference is better visualized in (C) and in a pseudocolor black and white image (D). (E-F) insets at higher magnification of images C and D. (B) Hoechst staining shows a clear difference in the nuclei staining in the posterior compartment that appears marked with a more intense staining due to the smaller size of the nuclei induced by the expression of *tor-RNAi*.

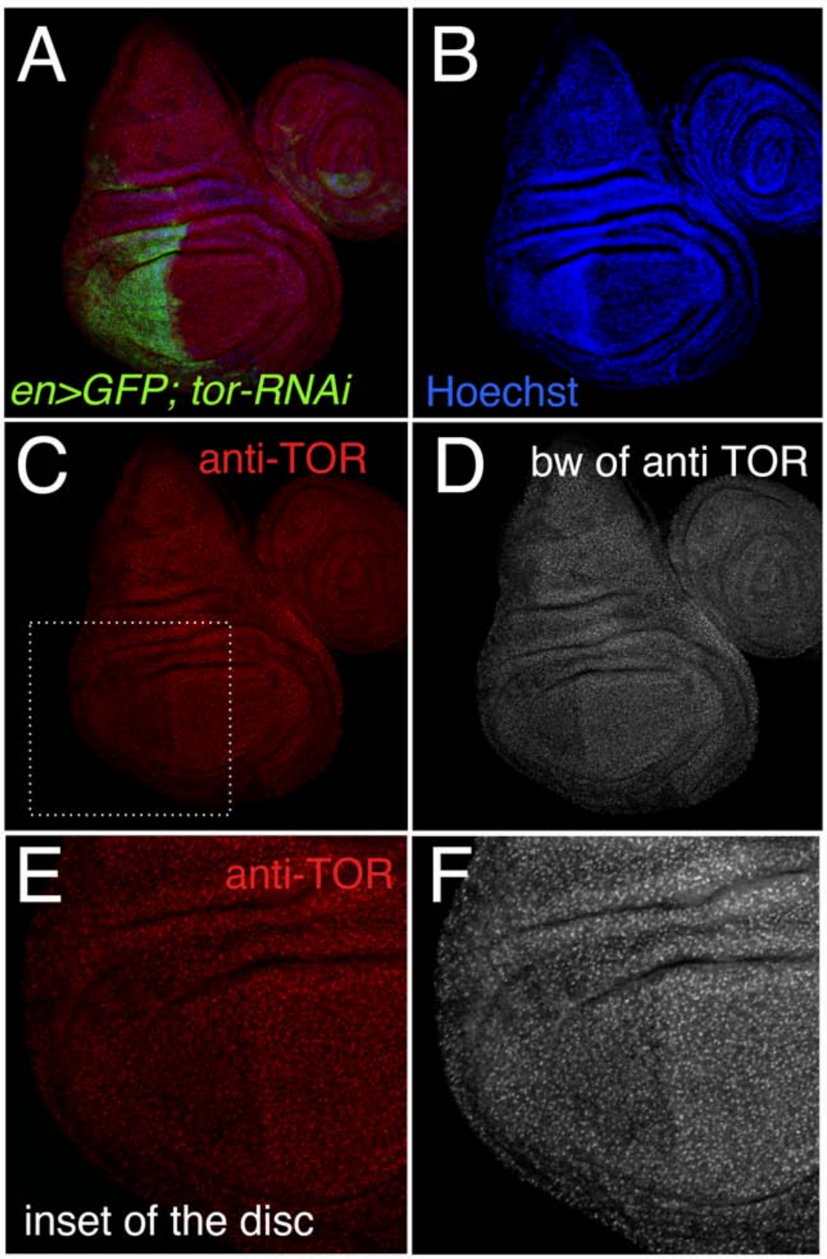

In addition, we also analyzed the ability of the antibody used for immune fluorescence in western blots using lysates from *tor-2L19 mutant* larvae [2] and control *w[1118]*. In these experiments the efficiency of our antibody was compared with an anti *Drosophila* d-TOR-P1 antibody developed by A. Teleman (DKFZ, Heidelberg) [3]. These experiments showed that the anti-mTOR antibody from ab45989 (B) recognizes a band of about 250 kDa (arrow), which is also recognized by an anti *Drosophila* TOR-P1 antibody (A) and only present in the lysates from *w[1118]* larvae and undetectable in lysates from *tor-2L19* mutant animals. Actin was used as control loading. Of note: ab45989 was raised against the 2150-2250 aminoacidic sequence of the human TOR protein (NP_004949.1) that shows 74% identity with *Drosophila* TOR protein (NP_001260427.1) from UniProt/BLAST program.

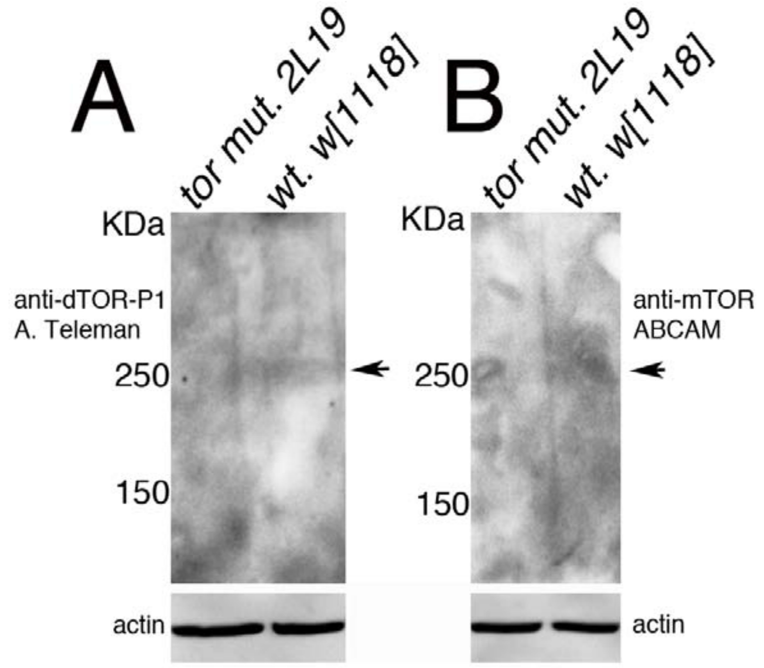


1. Torreira E, Seabra AR, Marriott H, Zhou M, Llorca O, Robinson CV, Carvalho HG, Fernandez-Tornero C, Pereira PJ: **The structures of cytosolic and plastid-located glutamine synthetases from Medicago truncatula reveal a common and dynamic architecture**. *Acta crystallographica Section D, Biological crystallography* 2014, **70**(Pt 4):981-993.

2. Oldham S, Montagne J, Radimerski T, Thomas G, Hafen E: **Genetic and biochemical characterization of dTOR, the Drosophila homolog of the target of rapamycin**. *Genes Dev* 2000, **14**(21):2689-2694.

3. Tsokanos FF, Albert MA, Demetriades C, Spirohn K, Boutros M, Teleman AA: **eIF4A inactivates TORC1 in response to amino acid starvation**. *The EMBO journal* 2016, **35**(10):1058-1076.
